# Effect of Highly Loaded Nanohydroxyapatite Composite Scaffolds Prepared via Melt Extrusion Additive Manufacturing on the Osteogenic Differentiation of Human Mesenchymal Stromal Cells

**DOI:** 10.1101/2021.01.21.427568

**Authors:** María Cámara-Torres, Ravi Sinha, Alberto Sanchez, Pamela Habibovic, Alessandro Patelli, Carlos Mota, Lorenzo Moroni

## Abstract

The field of bone tissue engineering seeks to mimic the bone extracellular matrix composition, balancing the organic and inorganic components. In this regard, additive manufacturing (AM) of highly loaded polymer-calcium phosphate (CaP) composites holds great promise towards the design of bioactive scaffolds. Yet, the biological performance of such scaffolds is still poorly characterized. In this study, melt extrusion AM (ME-AM) was used to fabricate poly(ethylene oxide terephthalate)/poly(butylene terephthalate) (PEOT/PBT)-nanohydroxyapatite (nHA) scaffolds with up to 45 wt% nHA, which presented significantly enhanced compressive mechanical properties, to evaluate their *in vitro* osteogenic potential as a function of nHA content. While osteogenic gene upregulation and matrix mineralization were observed on all scaffold types when cultured in osteogenic media, human mesenchymal stromal cells did not present an explicitly clear osteogenic phenotype, within the evaluated timeframe, in basic media cultures (i.e. without osteogenic factors). Yet, due to the adsorption of calcium and inorganic phosphate ions from cell culture media and simulated body fluid, the formation of a CaP layer was observed on PEOT/PBT-nHA 45 wt% scaffolds, which is hypothesized to account for their osteoinductivity in the long term *in vitro,* and osteoconductivity *in vivo*.

## 1. Introduction

Bone is one of the most recurrently damaged tissues in our body in form of fractures or defects caused by bone disease, traumatic injury, implant surgery, infection or tumor resection. [1] In case of long bones, bone loss in length can lead to critical-size defects, which will not heal on their own. Current strategies used in the clinics for the treatment of critical size bone defects mainly rely on i) distraction osteogenesis and induced membrane techniques, [2, 3] or ii) on the use of graft materials, such as autografts, allografts, or demineralized bone matrices, that fill and / or help to regenerate the defect. [4, 5] Treatment methods based on the first option are often complex, demanding for the patient, and require external fixation and multiple surgeries. Similarly, natural bone grafts, despite being osteoinductive and / or osteoconductive, possess several disadvantages, including donor site morbidity, restricted availability, high costs, pain, prolonged rehabilitation, lack of structural properties, and can trigger inflammation, immunological rejections or infections. [6] Therefore, the management of critical-size bone defects still remains a major clinical challenge, and consequently, a large amount of efforts are dedicated to investigate alternative strategies, such as tissue-engineered constructs to heal and regenerate bone in such scenarios. [7]

Mimicking the physical and chemical properties of bone has been the main approach towards the design of the optimal scaffold for bone regeneration. The bone matrix mainly consists of an organic phase of collagen type I, among other proteins, and an inorganic phase of carbonated hydroxyapatite. [8] Accordingly, bioceramics, and in particular calcium phosphates (CaP), have been extensively applied for hard tissue repair. Widely applied CaP phases include hydroxyapatite (HA), tricalcium phosphate (TCP), biphasic and triphasic calcium phosphate, and amorphous calcium phosphate, [9] and their osteoinductive and osteconductive performance is determined by various properties, such as chemical phase, structural properties (crystallinity, surface roughness, macro-, micro- and nanoporosity), mechanical properties and associated degradation behavior. [10] Yet, the role of the individual properties and their combination on the biological response to CaP-based bone graft substitutes has not yet been fully elucidated, with inconsistency in results found in the literature, which hampers the selection or design of the optimal CaP material for a specific application. [11, 12] Several reasons for this are that studies lack sufficient material and/or biological characterization, or use different cell types *in vitro*, unavoidable inter- and intra-species variations in *in vivo* experiments, convoluted material properties that cannot be easily tuned independently, and different fabrication techniques used for scaffolds manufacturing. [11, 12]

CaP processing into 3D macroporous scaffolds favors cell migration, tissue ingrowth, and vascularization. Moreover, the increase in surface area and body fluid accessibility provided by macroporosity can accelerate the degradation rate of the CaP material and boost the cell-material interactions towards an enhanced biological response, in terms of adhesion, proliferation and differentiation. [13] Compared to conventional 3D scaffold fabrication techniques for polymers, such as gas foaming, particulate leaching, freeze-drying and solvent casting, additive manufacturing (AM) has emerged as a promising macroporous scaffold fabrication technique. AM enables the fabrication of patient specific constructs with a fully interconnected network of pores, whose size and shape can be tuned to control cell response and the scaffold mechanical properties, of special importance for bone tissue engineering. [14] Nevertheless, pure CaP AM scaffolds, processed by powder bed and binder jetting techniques, require post-sintering steps, the removal of binders, and present poor mechanical properties, due to the inherent material brittleness and low fracture toughness, which overall limit their in vivo applications. [15, 16] Therefore, CaP have been processed by solution extrusion AM in combination with carrier liquids or solvents [17–19], or by melt extrusion AM in combination with synthetic biodegradable thermoplastic polymers. Among these, melt extrusion of polymer-CaP composite materials has been widely explored, as it mimics the organic-inorganic composition of bone tissue, displays a high processability window, and allows the fabrication of scaffolds with tunable degradation rates and mechanical properties. [20, 21] In particular, the combination of polymers with HA has gained interest, as HA closely mimics the bone mineral phase. Considering the average mineral content of bone tissue, which lies within the range of 50 wt% to 74 wt%, depending on the body location, [22, 23] tremendous efforts have been conveyed to increase the HA content in the scaffold, while retaining their printability and optimizing the mechanical properties. A few studies demonstrated feasibility of production of highly HA-loaded scaffolds processed by melt extrusion AM (ME-AM), including poly(ε-caprolactone) (PCL)-HA composite with 40 wt% ceramic phase, [24, 25] poly(lactic acid) (PLA)-HA composite with 50 wt% ceramic [26] and poly(ether urethane) (PEU)-HA composite with 40 wt% ceramic [27]. Despite material related advances, most of previously published reports focus only on the material synthesis and printability, and the structural characterization of the scaffold, [24–26, 28–30] while their biological properties are poorly characterized. [31, 32]

Here, we studied the *in vitro* osteogenic differentiation potential of highly loaded poly(ethylene oxide terephthalate)/poly(butylene terephthalate) (PEOT/PBT) – nano HA (nHA) composite scaffolds, fabricated by ME-AM. The biodegradable thermoplastic polyether☐ester multiblock copolymer PEOT/PBT was selected due to its processability, mechanical properties and proven *in vitro* and *in vivo* applicability in the orthopedic field. [33–37] While in previous studies this polymer has been blended with HA to a maximum of 25 wt% micro-HA, [38, 39] for the first time, PEOT/PBT-HA composite scaffolds with up to 45 wt% nHA were fabricated using ME-AM in the present report. To study their functionality in bone regeneration, human mesenchymal stromal cells (hMSCs) were chosen, due to the suitability of these stem cells over already committed cell lines to study the ability of a material to support osteogenic differentiation, [40] as well as for future clinical translation. We evaluated the effect of the scaffolds’ nHA content on the osteogenic differentiation of hMSCs by analyzing cell attachment, proliferation, matrix production and mineralization, and the expression of a relevant set of osteogenic genes and proteins.

## 2. Materials and methods

### 2.1. Polymer-nHA composite production by solvent blending and characterization

The production of PEOT/PBT-HA composites with 20 wt% and 45 wt% ceramic (20-nHA and 45-nHA, respectively) was carried out by a solvent blending procedure. PEOT/PBT pellets (PEO molecular weight = 300 kDa, PEOT:PBT weight ratio= 55:45, intrinsic viscosity 0.51 dl/g, Polyvation, The Netherlands) were dissolved in chloroform (Scharlab) under magnetic stirring for 45 min. An amount of 20 or 45 wt% nHA powder (particle size <200 nm, Sigma Aldrich) was added to the polymer solution and mechanically stirred at 750-1000 rpm for 15 min (Dissolver Dispermat CV3). After stirring, the dispersion was kept for 2h at 250 rpm, time during which part of the chloroform was evaporated and the viscosity of the blend increased. Subsequently, the polymer composite was precipitated in a mixture of diethylether and chloroform (Scharlab) (diethylether:chloroform = 10:1), and the supernatant removed. The composite was dried at RT, followed by a vacuum drying step at 90 °C for 2h to ensure the removal of solvent traces (diethylether <0.01 ppm, chloroform <1 ppm,). The foam-like composite formed during the precipitation step was compression molded at 150-175 °C and cut into pellets. The dispersion of nHA within the polymer was examined using backscattered scanning electron microscopy (BSEM, FEI’s Quanta 200 environmental scanning microscope, beam voltage 30 kV, spot size 5).

### 2.2. Scaffolds fabrication and characterization

45-nHA, 20-nHA and PEOT/PBT (0-nHA) scaffolds were fabricated via ME-AM. A commercial platform (Bioscaffolder 3.0, Gesim) was equipped with a custom-made printhead with two separate heating sources for the cartridge and extrusion screw. [41] The printing parameters applied to each material are briefly described in Table 1. A needle with 250 μm internal diameter was used to print 0-90 layout scaffolds, with 200 μm layer thickness and 750 μm strand distance (center to center). For cell culture experiments and scaffolds morphological evaluation, 4×15×15 mm^3^ (HxWxD) blocks were manufactured and cylindrical scaffolds of 4 mm diameter and 4 mm height were cored out using a biopsy punch. For mechanical testing, 4 mm diameter, 15 mm high cylindrical scaffolds were printed and directly used for analysis. These scaffold dimensions resemble the type of scaffold needed for a rabbit critical segmental bone defect study, as a real case example which will be performed in future experiments. The scaffolds’ architecture consisted also on 250 μm fiber diameter, 200 μm layer height, and 750 μm strand distance, and the scaffold layers had a 0-90 filament layout in the center, while at the edges, the filaments were deposited along the circular boundary. These scaffolds presented sufficient lateral porosity (Figure S4A-B).

**Table 1.** Printing parameters applied to obtained 3D scaffolds with each of the polymer composites

| Scaffold | T cartridge<br>(° C) | T screw<br>(° C) | Motor rotation<br>(rpm) | Cartridge Pressure<br>(bar) | Printing speed<br>(mm/s) |
| --- | --- | --- | --- | --- | --- |
| 0-nHA | 195 | 195 | 60 | 7 | 16 |
| 20-nHA | 205 | 210 | 45 | 8 | 18 |
| 45-nHA | 210 | 220 | 30 | 8 | 15 |

Scaffold morphology and porosity was assessed using a stereomicroscope (Nikon SMZ25) and scanning electron microscopy (SEM, XL-30, beam voltage 10 kV, spot size 3). Presence and distribution of fillers within gold sputter-coated scaffolds’ filaments surface and cross-section was examined using BSEM (XL-30, beam voltage 20 kV, spot size 5). SEM operating at 25 kV coupled with energy dispersive X-ray spectroscopy (EDS) was used to quantify the chemical composition of the samples. Particle size distribution on scaffolds was assessed on the acquired BSEM images and using the image analysis software tool XT Pro v3.2 (Soft Imaging System, GmbH).

Mechanical tests (compression at a rate of 0.1 mm/min., using a 100 N load cell) were carried out using an Instron Universal Mechanical test machine. Three scaffolds each, made from 0-nHA or 45-nHA were tested.

### 2.3. Cell seeding and culture

#### 2.3.1. Cell expansion

HMSCs isolated from bone marrow were purchased from Texas A&M Health Science Center, College of Medicine, Institute for Regenerative Medicine (Donor d8011L, female, age 22). Cryopreserved vials at passage 3 were plated at a density of 1,000 cells cm^−2^ in tissue culture flasks and expanded until approximately 80 % confluency in culture media (CM), consisting of αMEM with Glutamax, no nucleosides (Gibco), supplemented with 10% fetal bovine serum (FBS), without penicillin-streptomycin (PenStrep) at 37 °C and 5% CO_2_.

#### 2.3.2. Cell seeding in 3D scaffolds

Scaffolds were disinfected in 70% ethanol for 20 min, washed 3 times with Dulbecco’s PBS and incubated overnight in CM for protein attachment. Before seeding, scaffolds were dried on top of a sterile filter paper and placed in the wells of non-treated plates. Passage 4 hMSCs were trypsinized and resuspended in a dextran solution (10 wt/wt % dextran in CM, 500 kDa, Farmacosmos) to slow down cell settling and guarantee homogeneous cell attachment throughout the scaffold. Dextran is a temporary supplement and will be washed away during the first 24h. [42] A droplet of cell suspension (37 μl containing 200,000 cells) was placed on top of each scaffold before slowly filling the scaffolds’ pores. Seeded scaffolds were incubated for 4 hours at 37 °C and 5% CO_2_ to allow cells to attach. After this time, scaffolds were transferred to new wells containing 1.5 ml of basic media (BM), consisting of CM supplemented with 200 μM L-Ascorbic acid 2-phosphate and 1% PenStrep. Scaffolds were cultured for 7 days in BM followed by another 28 days in BM or mineralization media (MM), consisting of BM supplemented with dexamethasone (10 nM) and β-glycerophosphate (10 mM) (Sigma-Aldrich). Media was replaced after 24h and every two or three days during the culture period.

#### 2.3.3. Biochemical assays

##### 2.3.3.1. ALP activity and DNA assay

ALP activity and DNA content were evaluated over the culture period. For that, scaffolds were collected at every timepoint and washed with PBS, stored at −80 °C and freeze-thawed 3 times, to improve lysis efficiency. Samples were incubated for 1h at RT in a cell lysis phosphate buffer containing Triton X-100 (0.1 v %), at pH 7.8. After the addition of the chemiluminescent substrate for alkaline phosphatase CDP star® ready to use reagent (Roche) at a 1:4 ratio, luminescence was measured using a spectrophotometer (CLARIOstar®, BMG Labtech). Reported ALP values were normalized to DNA content, which was quantified using the CyQUANT cell proliferation assay kit (Thermo Fisher Scientific) on the remaining cell lysates. Samples for DNA quantification were incubated overnight at 56 °C in Proteinase K solution (1mg/ml Proteinase K (Sigma-Aldrich) in Tris/EDTA buffer) (1:1) for further cell lysis and matrix degradation. Subsequently, samples were freeze-thawed 3 times and incubated 1h at RT with a 20X diluted lysis buffer from the kit containing RNase A (1:500) to degrade cellular RNA. Samples were incubated with the fluorescent dye provided by the kit (1:1) for 15 min and fluorescence was measured (emission/excitation = 520/480 nm) with a spectrophotometer.

##### 2.3.3.2. ECM mineralization assay

Calcium deposits were qualitatively determined by alizarin red S staining (ARS) on composite scaffolds after 35 days of culture in BM or MM. Freshly fabricated scaffolds, and non-cell seeded scaffolds cultured in BM or MM and treated as cell seeded scaffolds in terms of media refreshment, were used as controls. Briefly, every timepoint scaffolds were fixed with 4% paraformaldehyde and washed with distilled water. Subsequently, scaffolds were stained with ARS (60 mM, pH 4.1 - 4.3) for 20 min at RT, and washed with distilled water. Scaffolds’ cross sections were imaged using a stereomicroscope. After imaging, ARS was extracted from the scaffolds by 1h incubation at RT with 1 ml 30 v% acetic acid while shaking, followed by 10 min at 85 °C. Afterwards, scaffolds were removed and solutions were centrifuged at 20,000 rcf for 10 min. An appropriate volume of ammonium hydroxide (5 M) was added to the supernatants to bring the pH to 4.2. The absorbance was measured at 405 nm using a spectrophotometer. Concentration of ARS was calculated from an ARS standard curve and the values were normalized to DNA content. The signal from control scaffolds was subtracted.

##### 2.3.3.3. Osteocalcin secretion

Osteocalcin secretion in media was analyzed after 14, 21 and 35 days of culture in BM or MM. Scaffolds’ culture media was refreshed on each timepoint by adding 1.5 ml of the corresponding media without FBS, in order to avoid background osteocalcin levels from the FBS. Upon collection (48 h later), media was centrifuged at 2,000 rcf for 10 minutes to remove debris and stored at - 80 °C until analysis. Osteocalcin content in medium was detected using a quantitative sandwich ELISA kit (ab195214) according to the manufacturer’s protocol.

#### 2.3.4. Gene expression

Gene expression analysis was performed after 7, 14, 21 and 35 days of culture. RNA was extracted from cells in scaffolds using a TRIzol isolation method. Briefly, TRIzol was added to each sample in an Eppendorf tube and centrifuged at 12,000 rcf for 5 min to precipitate the scaffold and ECM at the bottom. Chloroform was added to the supernatant and centrifuged at 12,000 rcf for 5 min to isolate the RNA, present in the aqueous phase. RNA was further purified using RNeasy mini kit column (Qiagen), according to the manufacturer’s protocol. The purity and quantity of total RNA was evaluated using a spectrophotometer. Reverse transcription was performed using iScript™ (Bio-Rad) following suppliers’ protocol. qPCR was performed on the mix composed of cDNA, SYBRGreen master mix (Qiagen) and the selected primers (Table 2) using a CFX Connect™ Real-Time System (Bio-Rad) under the following conditions: cDNA denaturation for 3 min at 95 °C, 40 cycles of 15 s at 95 °C, and 30 s at 65 °C. Additionally, a melting curve was generated for each reaction in order to test primer dimer formation and non-specific amplification. Gene transcription was normalized to the transcription of the housekeeping gene B2M. The 2^−ΔΔCt^ method was used to calculate relative gene expression for each target gene. Normalization was done with respect to relative gene expression of cells in 0-nHA scaffolds at day 7 in BM.

**Table 2.**
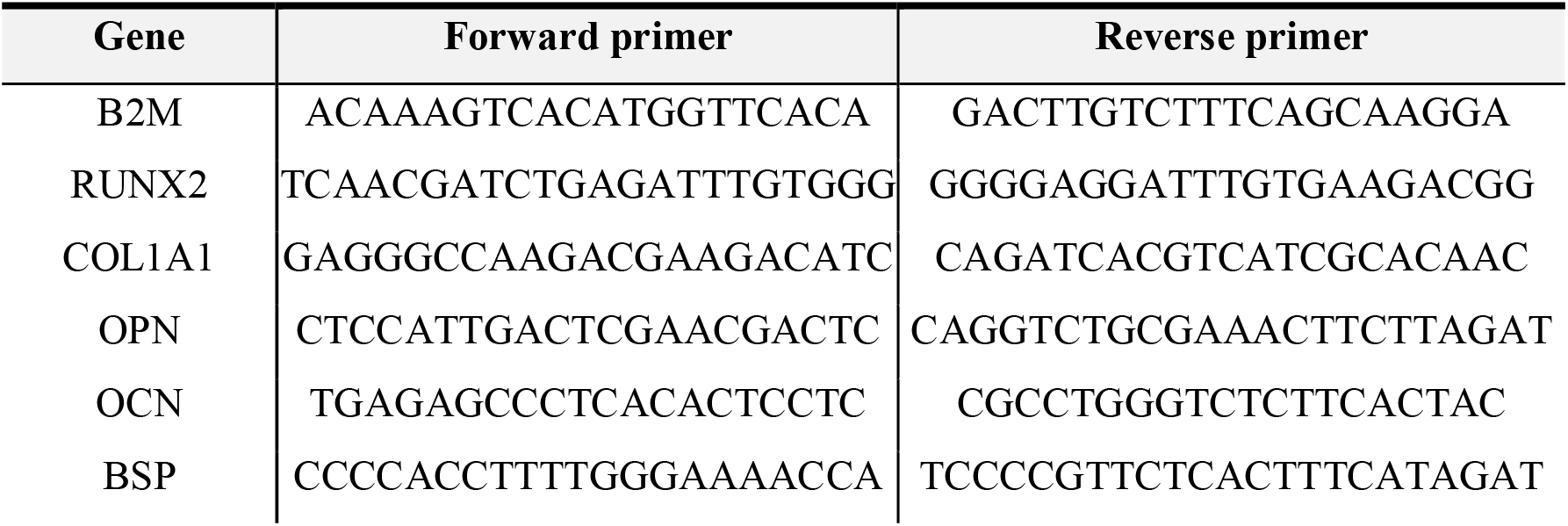
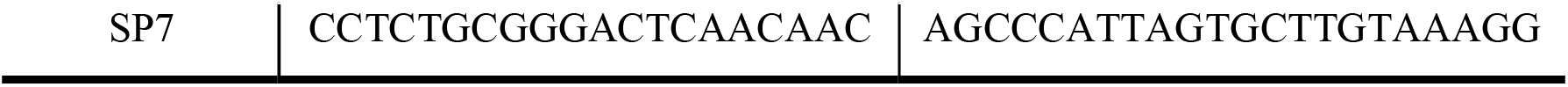
Primer sequences used for q-PCR. B2M: beta-2-microglobulin, RUNX2: runt-related transcription factor, COL1A1: collagen 1 alpha 1, OPN: osteopontin, OCN: osteocalcin, BSP: bone sialoprotein, SP7: osterix.

#### 2.3.5. Immunofluorescence

Cells in scaffolds were fixed in 4% paraformaldehyde after 35 days of culture in BM or MM, and permeabilized by 30 min incubation in Triton-X 100 (0.1 v%). Subsequently, the scaffolds were cut in half and washed 3 times with PBS. For assessing cell content and distribution on scaffolds, some scaffolds were directly stained for F-actin (488 Alexa Fluor Phalloidin, Thermo Fisher Scientific) for 1 h. Scaffolds cross-sections were imaged with a fluorescent microscope (Eclipse, Ti2-e, NIKON). Background subtraction and brightness adjustments were performed on the images using the software Image J, in order to improve their visualization. For antibody staining, scaffolds were further blocked for 1 h in blocking buffer (3 % BSA + 0.01% Triton-X 100). Afterwards, primary antibodies diluted 1:200 in washing buffer (10x diluted blocking buffer) were added (collagen I rabbit polyclonal (ab34710, Abcam), osteopontin rabbit polyclonal (ab8448, Abcam), osteocalcin rabbit polyclonal (ab93876)), each to a different sample half, and incubated overnight at 4 °C. Washed samples were incubated for 1h at RT with the secondary antibody (1:200 Alexa Fluor 647 goat derived anti-rabbit antibody, Thermo Fisher Scientific). Then, scaffolds were washed and stained for F-actin (488 Alexa Fluor Phalloidin, Thermo Fisher Scientific) for 1 h, followed by three washing steps with PBS. Confocal laser scanning microscopy was performed with a tandem confocal system (Leica TCS SP8 STED), equipped with a white light laser (WLL). Samples were excited with the dye specific wavelengths and emission was detected with HyD detectors. For optimal visualization in the reported images, F-actin was colored in green, collagen I in red, osteopontin in blue and osteocalcin in magenta.

#### 2.3.6. SEM, BSEM and EDS imaging of cultured scaffolds

After 35 days of culture, cell seeded and non-cell seeded scaffolds in BM or MM were fixed with 4 wt% paraformaldehyde for 30☐min. Scaffolds were dehydrated by a graded ethanol series (30, 50, 70, 80, 90, 96, 100 v%), during 15 min each at RT, followed by 15 min incubation in 100% ethanol-hexamethyldisilazane (HMDS) at a 1:1 ratio, and 15 min incubation in HMDS. HMDS was removed and scaffolds were allowed further drying overnight at RT in the fume hood. Samples were gold sputter coated. Cells, ECM and scaffolds surface were visualized on scaffolds cross-section using SEM (beam voltage 10 kV, spot size 3). The presence of mineral deposits on cell seeded scaffolds was examined by BSEM (beam voltage 20 kV, spot size 5). SEM operating at 25 kV coupled with EDS was used to identify Ca, P, and other elements on the ECM.

### 2.4. Bioactivity test in Simulated Body Fluid (SBF)

An SBF solution was prepared as previously described [43]. Briefly, chemicals were dissolved in distilled water in the following order: NaCl, NaHCO_3_, KCl, K_2_HPO_4_, MgCl_2_·6H2O, CaCl_2_·2H_2_O and Na_2_SO_4_. The solution was then buffered to pH 7.4 at 36.5° C using Tris and HCl (1M). The final solution had an ion concentration (mM) as follows: Na^+^, 142.0; K^+^, 5.0; Mg2^+^, 1.5; Ca2^+^, 2.5; Cl^−^, 147.8; (CO3)^−^, 4.2; (PO4)^2−^, 1.0; (SO4)^2−^, 0.5. Scaffolds were immersed in 1ml SBF and incubated at 37 °C. SBF was refreshed every 3-4 days. After 4 and 14 days, scaffolds were carefully rinsed in distilled water, blotted on an adsorbent paper and dried at RT. Then they were cut in half, gold sputtered and observed with SEM at 15 kV, spot size 3.

### 2.5. Ion release analysis

Before media refreshment at timepoints day 7, 14, 21 and 35 of the cell culture experiment, medium was collected from each scaffold and stored at −30☐°C until analysis. In a separated experiment, scaffolds were incubated in 1ml simulated physiological saline, which does not contain calcium or phosphate ions, (SPS; 0.8% NaCl, 50 mM HEPES, pH 7.3) for 28 days at 37 °C. This was done to analyze Ca and P release in a non Ca and P saturated solution. Every 3-4 days, an aliquot of the supernatant was collected, stored at −30☐°C until analysis, and the same volume of SPS was refreshed. Concentrations of Ca and P in cell culture medium and SPS were quantified by ICP-MS (iCaP Q, Thermo Scientific). Upon analysis, all samples, including fresh cell culture medium or fresh SPS, were thawed, vortexed for 30☐s, centrifuged at 20,000 rcf for 15 min to remove debris, and diluted 1:50 in 1 v% HNO_3_ with the addition of 20☐ppb scandium as internal standard. A standard curve of 20, 10, 5, 2.5 and 1.25 ppm Ca and P in 1 v% HNO_3_ and 20☐ppb scandium was prepared. Quantification was performed in STD mode with He as collision gas.

### 2.6. Statistical analysis

All data is shown as average with error bars indicating the standard deviation of at least three replicates. Analysis of statistics was conducted with GraphPad software. A one-way or two-way analysis of variance (ANOVA) was performed with a Tukey’s post-hoc multiple comparison test to evaluate statistical significance.

## 3. RESULTS

### 3.1. Scaffolds fabrication, nHA distribution and mechanical properties

Composite pellets prepared by solvent blending of PEOT/PBT and 20 or 45 wt% nHA (Figure S1) were used to fabricate PEOT/PBT-nHA composite scaffolds via ME-AM (Figure 1). SEM micrographs in Figure 1A,B showed that both 20-nHAand 45-nHA scaffolds maintained an interconnected macroporosity, comparable to the control 0-nHA scaffolds. BSEM images in Figure 1C and EDS analysis in Figure S2A confirmed an increasing amount of nHA, and therefore Ca and P, on the scaffold filaments with increasing nHA content in the composite blend. The presence of nHA on the surface of the 20-nHA and 45-nHA scaffolds was further confirmed by ARS, which binds to the calcium of nHA (Figure S2B, C). An increase in the pink-red stain intensity on the surface of scaffolds was observed with increasing nHA content. Despite the nanometer size of the raw nHA (<200 nm), particle aggregation was already noticed in the solvent blending process prior to melt extrusion, possibly occurring during the precipitation step of the polymer composite solution in the non-solvent (Figure S1B). In the 45-nHA scaffolds, nHA micro aggregates occupied around 24% of the filaments’ surface area. Quantification on their size distribution revealed that ~75% of the nHA micro aggregates possessed a diameter in the range of 2-10 μm, while ~5% of the particles diameter lied in the range of 30 to 90 μm (Figure S3). The other 76% of the 45-nHA scaffolds filaments surface area was occupied by the polymer and nanometer size nHA particles. The presence of these nHA micro aggregates provided a higher surface roughness to the composite scaffolds when compared to the smooth 0-nHA scaffolds (Figure 1B). The addition of nHA up to 45 wt% led to a 2-fold increase in the scaffolds’ compressive modulus, compared to copolymer only scaffolds (Figure S4C). Compressive strength at yield was also slightly increased on the 45-nHA scaffolds (Figure S4D). Yet, no significant differences were observed compared to 0-nHA scaffolds.

**Figure 1.**
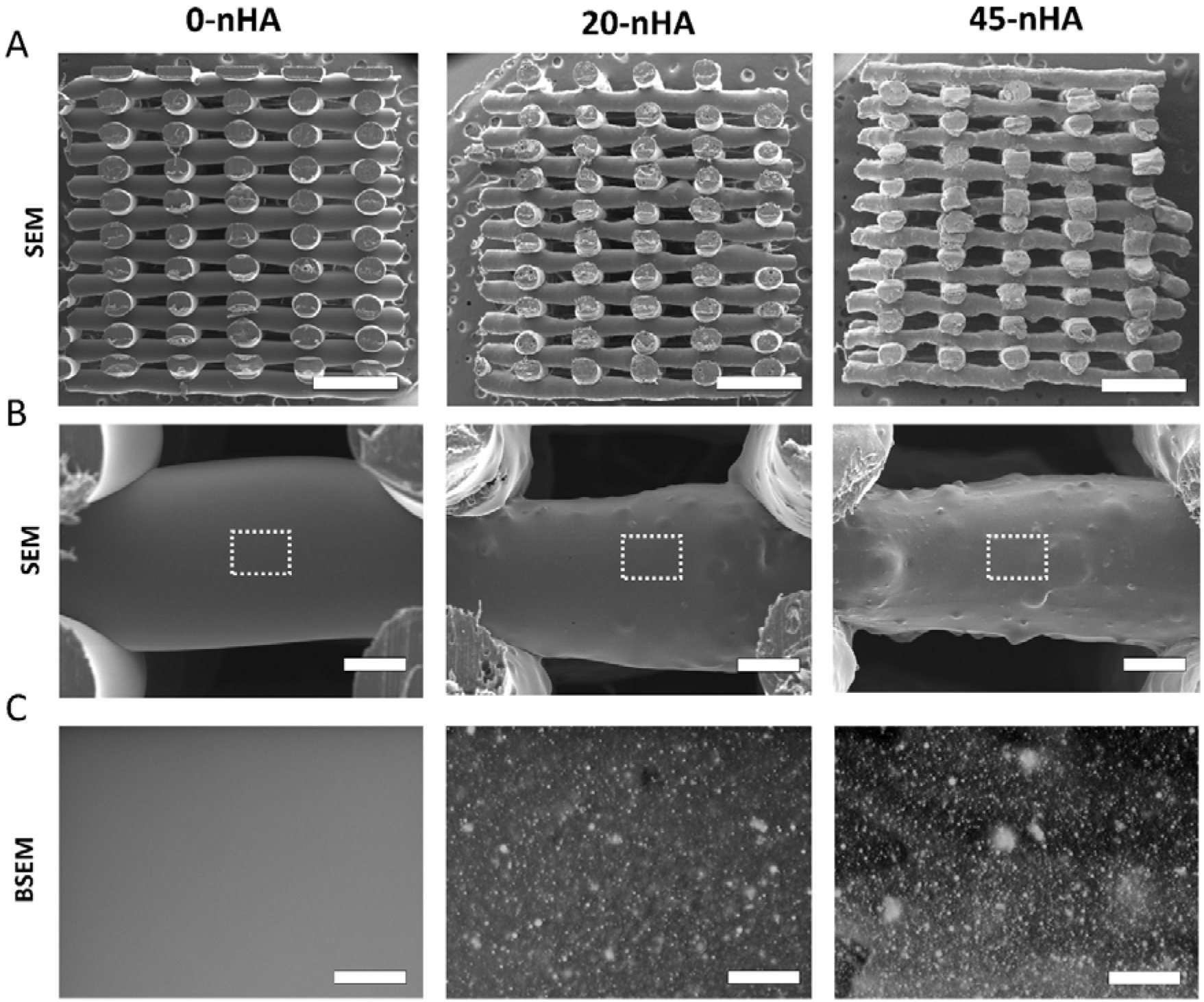
PEOT/PBT-nHA composite scaffolds fabrication and nHA distribution in scaffold filaments. (A) SEM micrographs of scaffolds cross-section showing their macroporosity. Scale bars: 1 mm. (B) Representative BSEM micrographs of scaffolds filaments depicting nHA distribution and increasing surface roughness with increasing nHA content. Scale bars: 100 μm. The regions marked with dashed lines are magnified in (C). Scale bars: 20 μm.

### 3.2. hMSCs seeding efficiency, proliferation and osteogenic differentiation on PEOT/PBT-nHA composite scaffolds

Scaffolds were seeded using a viscous seeding solution to ensure homogeneous hMSCs attachment throughout the scaffold. Consequently, all scaffolds qualitatively showed uniform cell distributions from the earliest timepoints. Moreover, high seeding efficiency values in the order of ~80 % were maintained among different scaffold types without statistical differences in terms of cell attachment (Figure 2A). To understand the effect of nHA on the osteogenic differentiation of hMSCs, scaffolds were cultured up to 35 days in BM or MM. To support cell proliferation before the addition of osteogenic factors, scaffolds were cultured initially for 7 days in BM. Yet, DNA content analysis demonstrated no cell proliferation during this period (Figure 2A). DNA content was further monitored over the culture time (Figure 2B). Although no statistical differences were observed in terms of cell number among scaffold types and time points, a decrease in cell number from day 14 to day 35 of culture in BM was observed for all scaffold types. Overall, Figure 2B also demonstrated lower cell numbers in scaffolds cultured in BM when compared to MM at late timepoints (day 21 and day 35).

**Figure 2.**
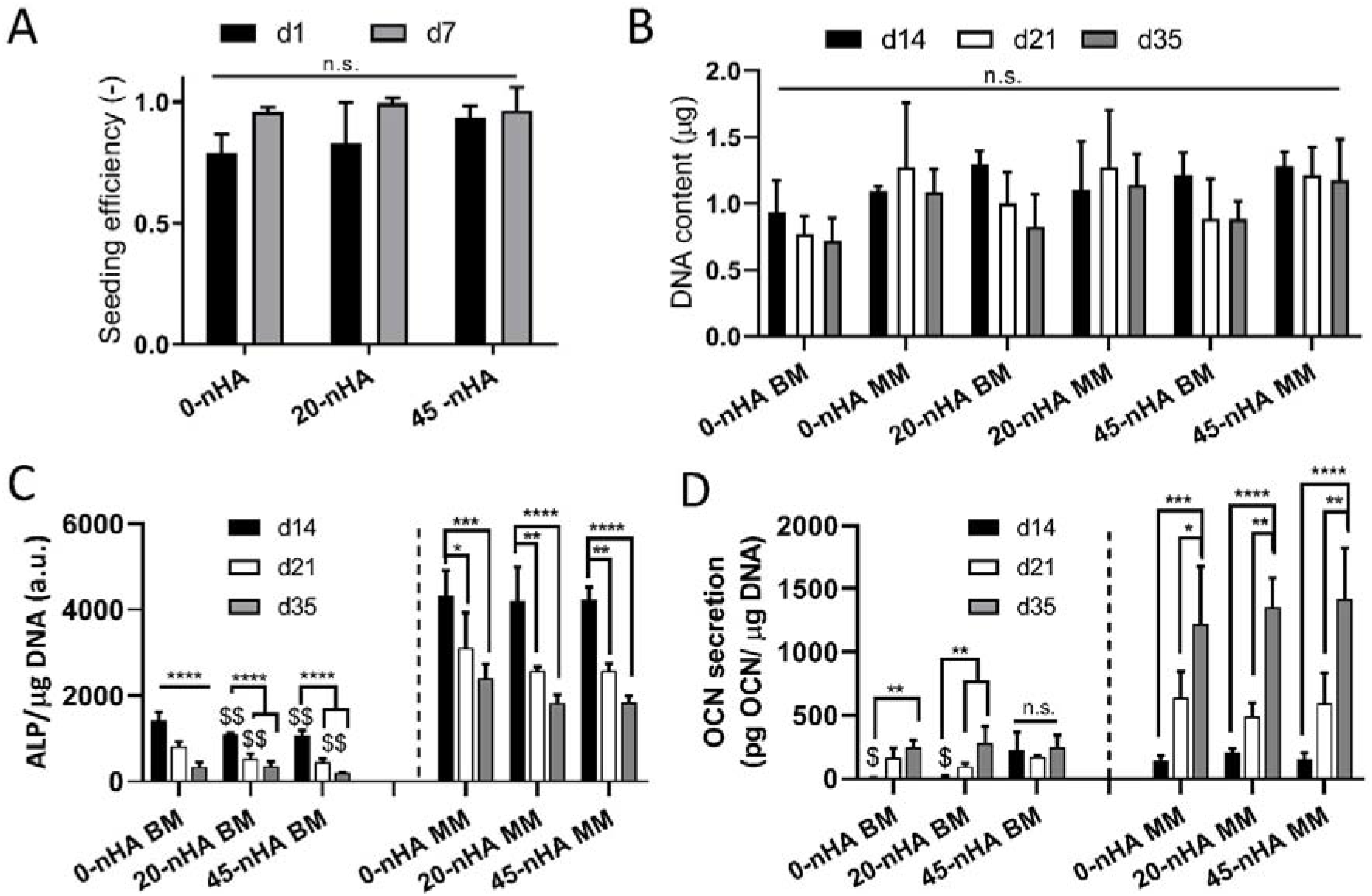
Cell seeding and osteogenic differentiation on PEOT/PBT-nHA composite scaffolds. (A) Seeding efficiency and proliferation 7 days after seeding. (B) DNA content progression on 0-nHA, 20-nHA, 45-nHA scaffolds over 35 days of culture in BM or MM. (D) ALP activity of hMSCs progression over 35 days when seeded on the different scaffold types in BM or MM. (E) OCN secretion by hMSCs in media progression over 35 days when seeded on the different scaffold types in BM or MM. Data presented as average ± s.d. and statistical significance performed using two-way ANOVA with Tukey’s multiple comparison test. For (A,B): n.s. p > 0.5. For (C): *$ p < 0.05; **$$ p < 0.001; ***$$$ p < 0.001; ****$$$$ p < 0.0001; * for comparisons among timepoints each scaffold type; $ for comparisons to 0-nHA each timepoint; comparisons among BM and MM samples performed separately. For (D): n.s. p > 0.05; *$ p < 0.05; **$$ p < 0.001; ***$$$ p < 0.001; ****$$$$ p < 0.0001; * for comparisons among timepoints each scaffold type; $ for comparisons to 45-nHA each timepoint; comparisons among BM and MM samples performed separately.

ALP activity, an early osteogenic marker, was evaluated at day 14, 21 and 35 (Figure 2C). ALP values were higher when cells were cultured in MM compared to BM. A peak in ALP activity was shown at day 14, with a progressive and significant decrease on day 21 and day 35, both in BM and MM. While in MM no statistical differences among scaffold types were observed at any time point, ALP values in BM were found to be significantly higher at day 14 and 21 in 0-nHA scaffolds compared to 20-nHA and 45-nHA scaffolds. When analyzing osteocalcin (OCN) secretion in media, higher values were observed in MM than in BM (Figure 2D). Since OCN is a late marker for osteogenic differentiation, a peak was observed at day 35, in contrast to ALP. As noted with ALP, no significant differences were observed in OCN secretion among scaffold types in MM. Nevertheless, a significantly higher OCN signal was recorded on 45-nHA scaffolds already at an early time point (day 14) when cultured in BM, compared to 20-nHA and 0-nHA scaffolds in BM. Notably, other scaffold types cultured in BM reached this levels of OCN secretion at day 21 or 35 at the earliest.

#### 3.2.1. Gene expression

The expression of bone related transcription factors and protein encoding genes was assessed at day 14, 21 and 35, under BM or MM conditions. Figure 3 shows the relative gene expression change with respect to 0-nHA day 7 BM condition (Figure S5). In general, RUNX2 gene expression was found to be higher in MM than in BM for all scaffold types over the whole culture period (Figure 3A). At day 14 in MM, significantly higher expression on 0-nHA scaffolds was observed. However, no significant differences among scaffold types were observed at later time points in MM. Similarly, despite higher COL I gene expression on 0-nHA scaffolds at day 14 in MM compared to the nHA composite scaffolds, comparable COL I gene expression levels were maintained at later culture time points among all scaffold types (Figure 3B). Interestingly, the culture media appeared not to have any effect on COL I gene expression and no significant differences were found between BM and MM at a given time point and scaffold type over the culture period. In the case of OPN gene expression, higher levels were noticed when cultured in BM compared to MM for all scaffold types at days 14 and 21 (Figure 3C). Yet, at day 35 the gene expression level of OPN was comparable among all scaffold types in both BM and MM media. Independent of the culture media, a low gene expression of OCN was maintained in all scaffold types until a significant upregulation, slightly higher in 0-nHA scaffolds, occurred at day 35 in MM (Figure 3D). A significant upregulation in the BSP and Osterix gene expression was observed for all scaffold types cultured in MM from day 14 to day 21, and to day 35 in the case of Osterix (Figure 3E, F). At day 35, gene expression levels in 0-nHA and 45-nHA remained comparable and significantly higher than in BM.

**Figure 3.**
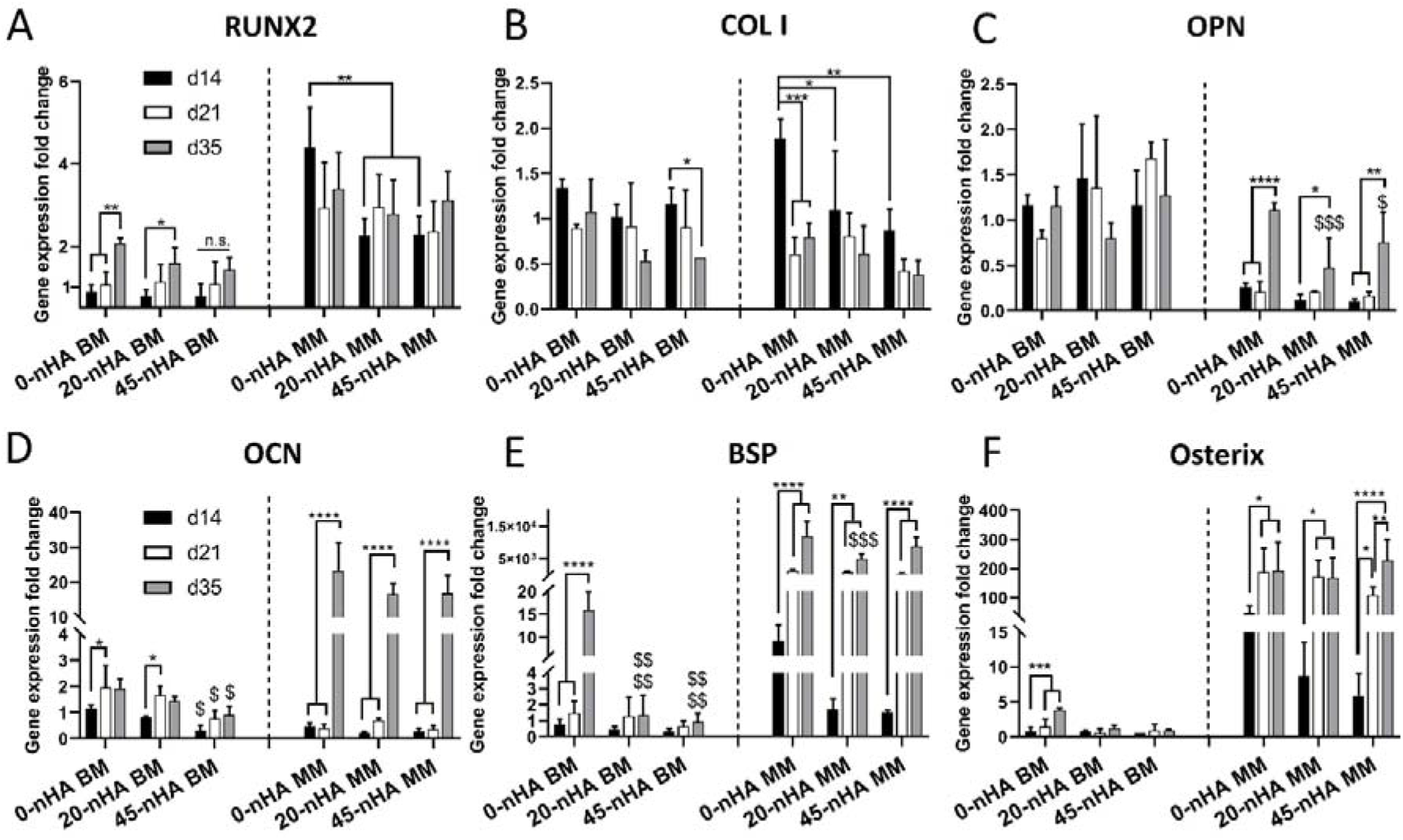
Gene expression of hMSCs cultured on scaffolds with different nHA concentrations for 14, 21 and 35 days in BM or MM. (A) RUNX2, (B) COL 1, (C) OPN, (D) OCN, (E) BSP, and (F) Osterix fold change expression values relative to 0-nHA d7 BM. Data presented as average ± s.d., and statistical significance performed using two-way ANOVA with Tukey’s multiple comparison test (*$ p < 0.05; **$$ p < 0.01; ***$$$ p < 0.001; ****$$$$ p < 0.0001; * for comparisons among timepoints each scaffold type; $ for comparisons to 0-nHA (control) each timepoint; comparisons among BM and MM samples performed separately).

#### 3.2.2. Protein expression

Complementary to gene expression, protein expression at day 35 was evaluated by immunofluorescence on all scaffold types and culture conditions. Extracellular COL1 was found covering filaments on all scaffolds types and culture media conditions (Figure 4). Supporting the gene expression profiles, OPN protein expression was found slightly more abundant on scaffolds cultured in BM compared to MM, except in 45-nHA scaffolds in MM, where OPN protein expression was higher than in 20-nHA or 0-nHA scaffolds. Following an opposite trend, OCN protein expression was higher in scaffolds cultured in MM compared to BM (Figure 4). Interestingly, OCN protein expression was also recorded on nHA containing scaffolds (20-nHA and 45-nHA) cultured in BM.

**Figure 4.**
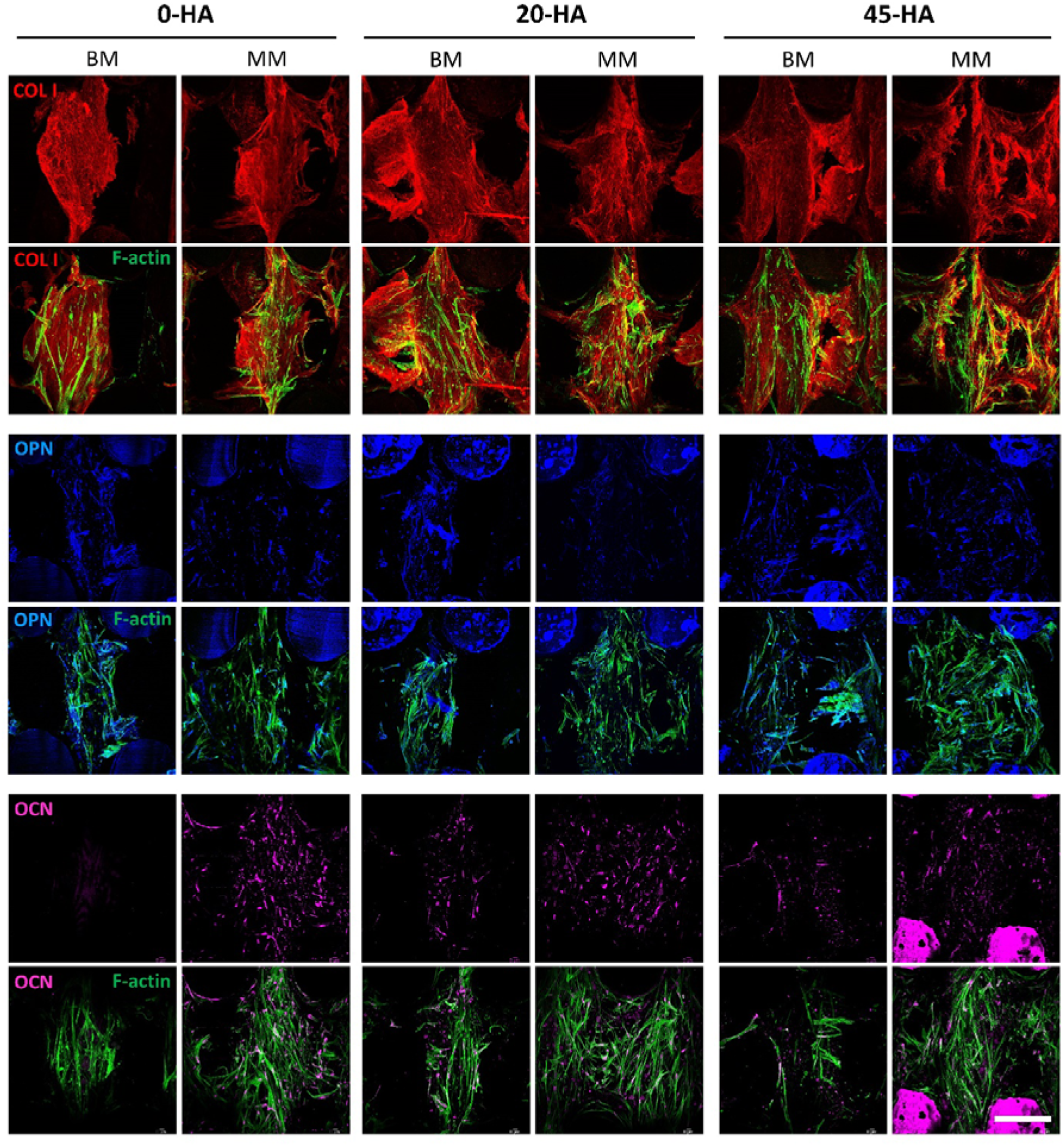
Influence of nHA content on the hMSCs expression of different osteogenic proteins. Representative confocal microscopy images of hMSCS (F-actin, green) on top of the filaments of 0-nHA, 20-nHA and 45-nHA scaffolds after 35 days of culture in BM or MM and stained for the relevant osteogenic markers COL I (red), OPN (blue) and OCN (magenta). Scale bar for all images: 200 μm.

#### 3.2.3. Extracellular matrix (ECM) production and mineralization

ECM mineralization was examined at day 35 of culture in BM and MM. An increasing amount of calcium deposition with increasing nHA content was observed, both quantitatively (Figure S6) and qualitatively, in scaffolds cultured in MM (Figure 5A, Figure S7). Although at lower levels than in MM, ARS quantification also revealed calcium deposits on 20-nHA and 45-nHA scaffolds when cultured in BM (Figure S6), after subtraction of the signal given by non-cell seeded stained scaffolds. In BM calcium deposition was mostly appreciable on 45-nHA scaffolds, where a more intense red color, compared to the non-seeded controls, was observed covering most of the surface area of the 45n-HA scaffolds filaments, as well as the scaffold filaments’ exposed cross sections on the scaffolds outer surface (Figure 6A, Figure S7).

**Figure 5.**
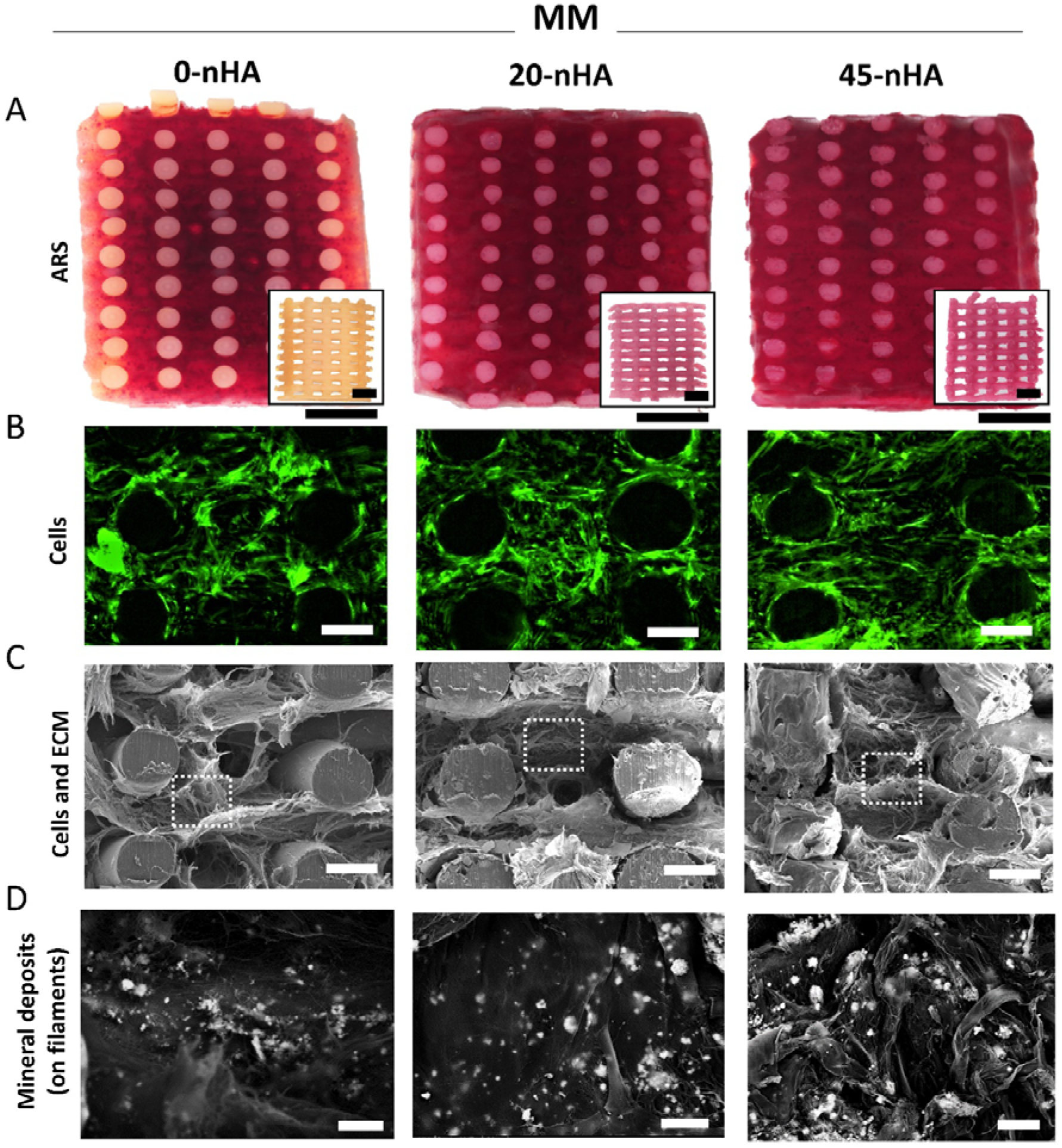
Influence of scaffolds’ nHA content on matrix mineralization when cultured for 35 days in MM (7 days in BM followed by 28 days in MM). (A) Representative stereomicroscope images of scaffolds cross sections stained with Alizarin Red S (ARS). Inserts represent the corresponding control scaffolds without cells incubated in MM and stained with ARS. Scale bars: 1 mm. (B) Representative fluorescent microscopy images of hMSCs (F-actin, green) on scaffolds’ cross sections. Scale bars: 200 μm. (C) Representative SEM micrographs of scaffolds’ cross sections depicting cell and ECM coverage. Scale bars 200 μm. The regions marked with dash lines (ECM in scaffolds’ pores) are magnified and visualized in BSEM modality in (D) for examining the mineral deposits. Scale bars: 20 μm.

**Figure 6.**
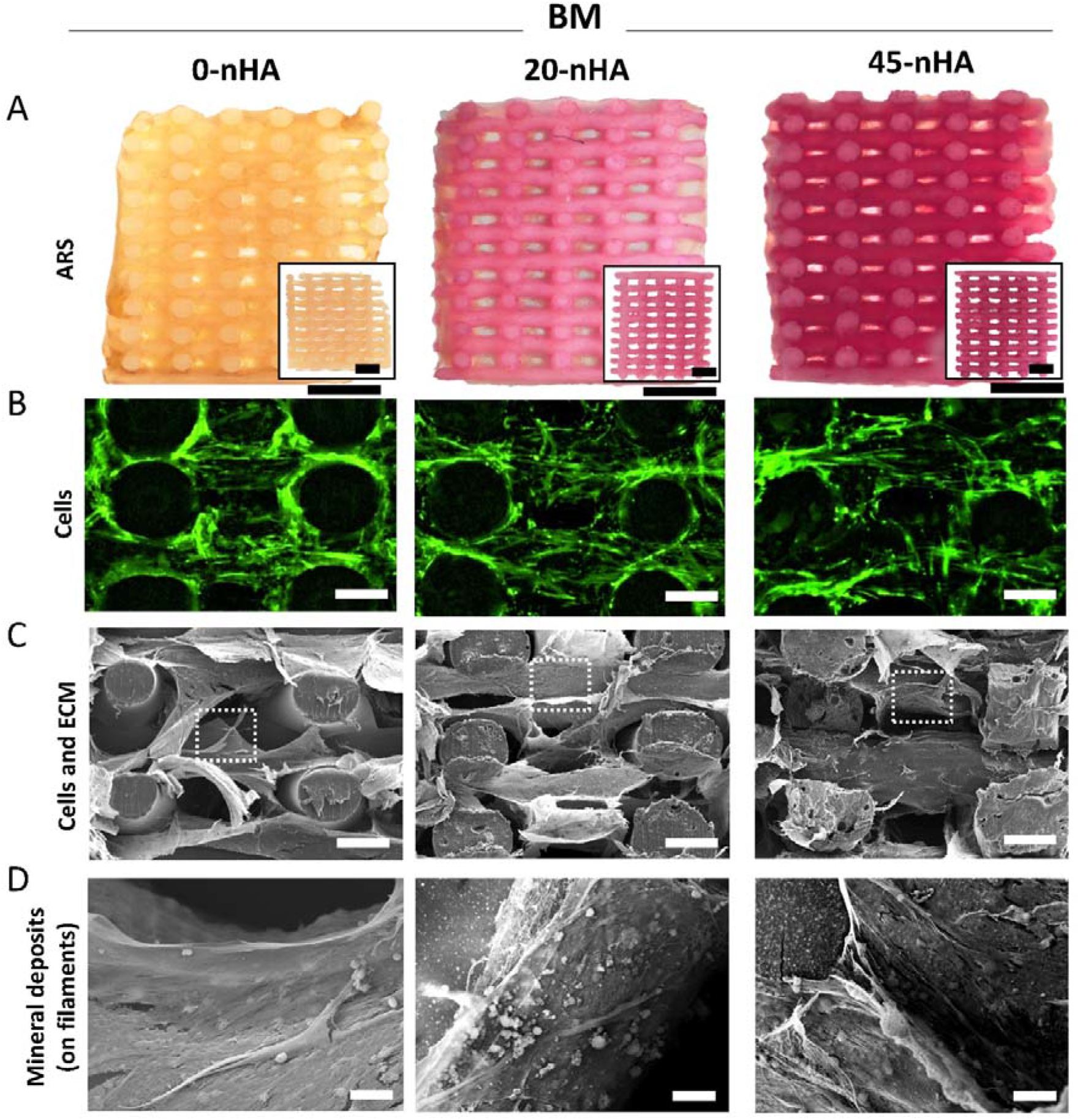
Influence of scaffolds’ nHA content on matrix mineralization when cultured for 35 days in BM. (A) Representative stereomicroscope images of scaffolds cross sections stained with Alizarin Red S (ARS). Inserts represent the corresponding control scaffolds without cells incubated in BM and stained with ARS. Scale bars: 1 mm. (B) Representative fluorescent microscopy images of hMSCs (F-actin, green) on scaffolds’ cross sections. Scale bars: 200 μm. (C) Representative SEM micrographs of scaffolds’ cross sections depicting ECM coverage. Scale bars: 200 μm. The regions marked with dash lines (ECM in scaffolds’ filaments) are magnified and visualized in BSEM modality in (D) for examining the mineral deposits. Scale bars: 20 μm.

Next, cell and ECM coverage at day 35 of culture was investigated. When cultured in MM, cells in all scaffold types were confluent occupying both the scaffolds’ filaments and pores along the whole scaffold cross-section (Figure 5B and Figure S8). Moreover, a dense and fibrillar ECM network produced by cells was observed for all scaffold types (Figure 5C and Figure S9). BSEM images in Figure 5D revealed bright points intercalated within the ECM in scaffolds pores, which were discriminated via EDS as Ca and P deposits, as well as a minimal amount of Na salts from the culture medium (Figure S10A). In addition, a peak of N, corresponding to the ECM proteins, was also observed. In contrast to the dense ECM in scaffolds cultured in MM, a thin layer of cells and ECM was visualized covering the scaffolds’ filaments when cultured in BM (Figure 6B-C, Figure S8, Figure S9). Notably, BSEM images of the ECM of cells cultured in BM did not depict bright points corresponding to mineral deposits (Figure 6D) and EDS analysis demonstrated significantly lower Ca and P signals compared to the analysis on scaffolds cultured on MM (Figure S10B). To be able to better discern among the EDS signal coming from the Ca and P already contained in the scaffold material, and the newly deposited minerals, not only the cell and ECM directly deposited on top of the scaffolds filaments was analyzed, but also the cell and ECM layer detached from the filament or present in the pores. Ca and P content in the ECM on the pores of MM scaffolds was found to be significantly higher than that on BM scaffolds (Figure S10).

### 3.3. Ca and P ion exchange dynamics with the medium

To investigate the degradation of the nHA contained in the scaffolds, Ca release into SPS, which does not contain these ions, was quantified by ICP-MS. nHA containing scaffolds released Ca into SPS continuously over the period evaluated (26 days) (Figure S11). In agreement with the nHA content on the scaffolds, 45-nHA scaffolds released larger amounts and at a higher rate (50 - 200 μM every 3-4 days) than 20-nHA scaffolds (10 - 50 μM every 3-4 days).

To investigate the Ca and P content of the medium when scaffolds are in a cell culture setting, ICP-MS analysis was carried out on media collected at different time points during culture. This enabled to elucidate the ion exchange dynamics of Ca and P between the scaffolds and the media. The lower concentrations of Ca in media in contact with scaffolds compared to fresh media concentrations demonstrated that all scaffolds, cell seeded and non-cell seeded, depleted the media from Ca, especially when cultured in MM (Figure S12). Similarly, scaffolds cultured in MM depleted the media from P, while P levels in BM remained similar to fresh media ones. Interestingly, in general no significant differences in Ca and P depletion from media were found among cell seeded and non-cell seeded control scaffolds, nor among scaffolds with different nHA concentrations (Figure S12). To complement these results, the surface of the filaments of cell seeded (areas without cells) and non-cell seeded scaffolds cultured in BM and MM were analyzed by SEM and EDS (Figure S13 and Figure S14). Interestingly, while according to ICP-MS 0-nHA scaffolds depleted the media from Ca in BM, and from Ca and P in MM, no Ca and P was observed on any of the 0-nHA scaffolds surface. In contrast, the surface of 20-nHA and 45-nHA scaffolds in MM was found to be covered by CaP spheres in non-cell seeded scaffolds and by a compact CaP layer in cell seeded scaffolds. In BM, these scaffolds showed a higher amount of white particles in their surface, compared to as-prepared scaffolds, potentially corresponding to the deposition of a Ca based mineral phase. While not easy to verify either by imaging or EDS, the surface of 45-nHA scaffolds in BM also looked more populated with bright spots, potentially corresponding to Ca deposits.

The scaffolds were also immersed in SBF, a solution that closely mimics human blood plasma and is used to test the ability of a material to deposit mineral on its surface. SEM showed a more pronounced deposition of mineral on 45-nHA scaffolds compared to 20-nHA scaffolds (Figure 7). As expected, no mineral deposition occurred on the 0-nHA scaffolds even after 14 days of immersion, when a thick CaP layer was already covering the 45-nHA scaffolds.

**Figure 7.**
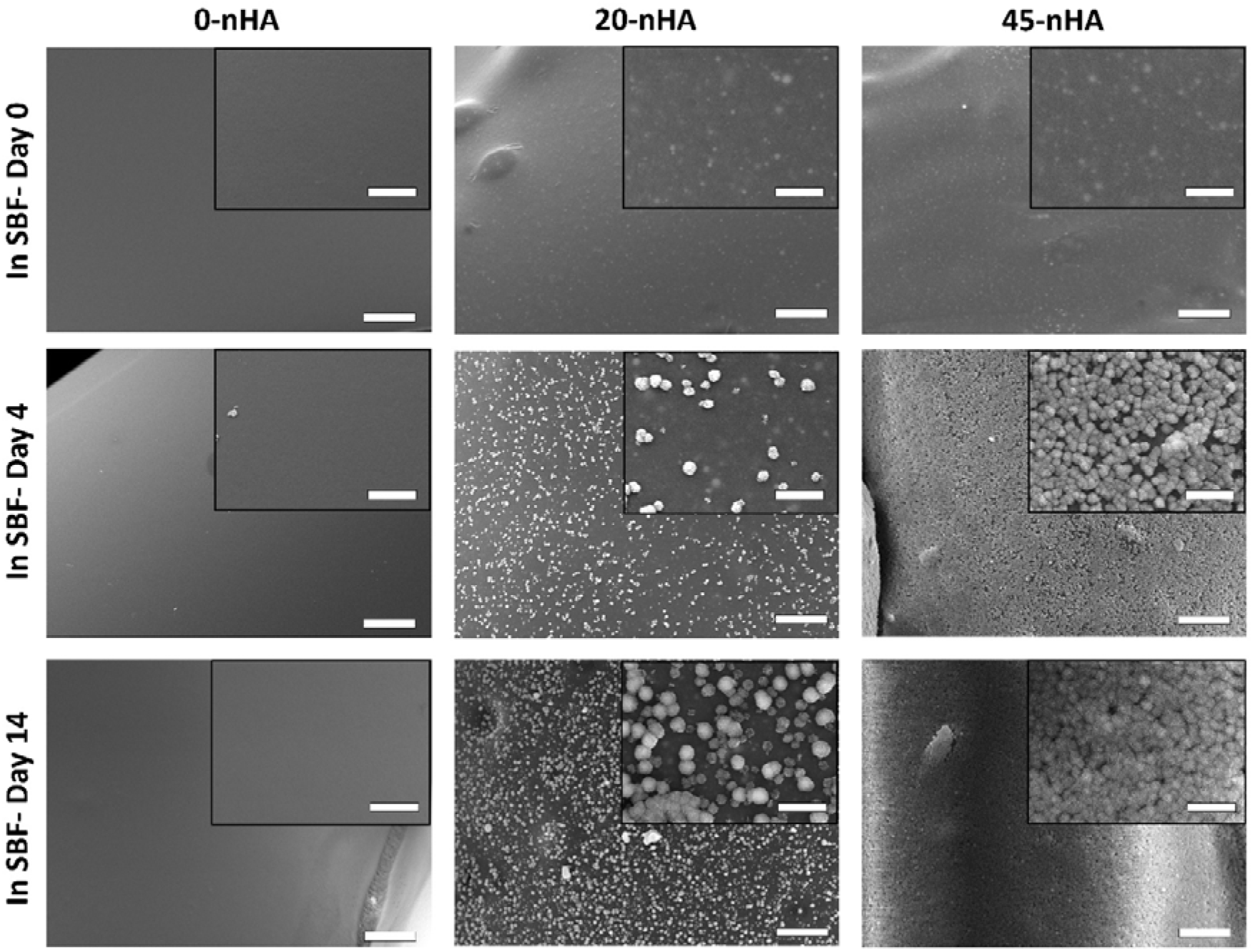
Representative SEM micrographs of scaffolds’ filaments surface mineralization after immersion in SBF for 4 and 14 days. As-prepared scaffolds (day 0) are presented as controls. Scale bars: 20 μm. Inserts represent magnified regions. Scale bars 5 μm.

## 4. DISCUSSION

During the last decades, AM has rapidly grown into the field of tissue engineering offering patient specific 3D scaffolds from a wide range of biocompatible materials, including polymer-CaP composites. CaP fillers are highly desired when aiming to mimic the composition of bone and to provide scaffolds with high strength, tunable degradation and favorable bioactivity. While solvent extrusion has permitted the fabrication of AM scaffolds with up to 90 wt% HA, [44, 45] the need of low volatility organic solvents, i.e. plasticizers, has shown to lead to solvent remnants within the scaffold, as well as to elastic constructs with poor applicability in load bearing applications. The advantage of thermoplastic ME-AM for bone tissue engineering over other AM techniques, such as stereolithography, mainly lies in the possibility of processing viscous composite materials including fillers, such as CaP particles, in a cost effective manner, and without requiring the use of organic solvents. Since the firstly reported PCL-CaP ME-AM scaffolds, containing up to 25 wt% CaP, [46–49] the field has largely evolved seeking to fabricate scaffolds with an inorganic content closer to native bone ECM, and with new polymeric materials. Here, we were able to fabricate composite 3D scaffolds based on PEOT/PBT, which have a great potential in the orthopedics field, and with up to 45 wt% nHA, closer to native bone mineral content than the majority of the until now reported scaffolds. Despite the high nHA loading, composites were printable and scaffolds’ filaments maintained a good shape fidelity and macroporosity (pore size 500 μm), which is expected to favor tissue infiltration and vascularization *in vivo*. [50] When looking at the filaments’ surface and cross-sections of the composite scaffolds, nHA aggregates with a size in the order of tens of microns were observed, in accordance with previous reports, where melt extruded filaments with HA concentrations above 10 wt% demonstrated HA agglomeration. [51] In spite of these microaggregates, nanosized particles were still present in a well-dispersed manner within the polymer matrix of the scaffolds and on their surface, which were expected to elicit a higher degree of biological response compared to microsized counterparts, due to their high surface-to volume-ratio and to their small size, allowing uptake by cells. [52, 53] Most importantly, the high nHA content significantly enhanced the mechanical properties of the scaffolds, with 45-nHA scaffolds having a compressive modulus of 92 ± 25 MPa and compressive strength of 4.4 ± 0.9 MPa, both in the range of cortical bone mechanical properties. [54, 55] Moreover, these scaffolds are stiffer than previously reported melt extruded AM scaffolds prepared with PCL-HA composites with similar inorganic loadings. This suggests the suitability of 45-nHA scaffolds for their application in bone regeneration scenarios, compared to bare PEOT/PBT scaffolds, as well as highly loaded PCL scaffolds, which is a commonly used polymer for bone scaffolds. [24, 25]

*In vivo*, osteoinductive materials have the ability of recruiting stem cells and differentiating them towards an osteogenic phenotype in a heterotopic location. [12, 56] Due to the complexity of the *in vivo* microenvironment, *in vitro* models do not always reliably predict osteoinductivity. [57] Yet, culturing stem cells on biomaterial scaffolds and analyzing cells’ ability to differentiate can give some initial insights into a material osteogenic potential and aid to understand the osteoinduction mechanism. Thus, we cultured hMSCs on scaffolds in BM, without osteogenic factors, i.e. dexamethasone and beta glycerol phosphate, and assessed their phenotype as a function of nHA content. In addition, we evaluated the synergistic effect of the material and these soluble factors that stimulate osteogenic differentiation/mineralization, by culturing in MM. Previously, it has been shown that cell distribution and density upon seeding can significantly influence hMSCs osteogenic differentiation in 3D scaffolds. [42, 58, 59] Accordingly, poor cell attachment and lack of confluency within the scaffold cross-section have been shown to result in poor differentiation and lack of matrix mineralization *in vitro.* Besides, some studies reported enhanced cell attachment on polymer-CaP composites, likely due to the hydrophilicity and protein adsorption capacity of the CaP [60, 61]. In the present study, in order to decouple the osteogenic potential of the PEOT/PBT-nHA composite scaffolds from the effect of attachment efficiency, a viscous solution was used as seeding media, thereby ensuring comparable cell attachment and homogeneity among all scaffolds regardless of the nHA content, as previously reported. [42] Due to scaffold surface saturation with cells upon seeding, no cell proliferation was observed in the first 7 days of culture, nor during the subsequent 28 days in BM on any scaffold type, as ECM production was limited to the scaffolds filaments surface. On the other hand, ECM was also produced within the pores when cultured in MM, which increased the growth surface area enabling cell migration and a slight, yet not statistically significantly different, increase of cell number. An important general conclusion that can be drawn from the DNA quantification, and cell and ECM imaging on the scaffolds, is that 3D ME-AM PEOT/PBT scaffolds with up to 45 wt% nHA are able to support cell growth, both in BM and MM.

Osteogenic differentiation was initially evaluated by ALP activity, an early marker for osteogenesis, [62] and osteocalcin secretion, one of the major bone non-collagenous proteins with the ability to bind to the bone HA. [63] Moreover, the gene expression of RUNX2, an essential transcription factor to boost the expression of bone ECM proteins [64, 65], as well as other bone markers for osteogenesis at mRNA level (i.e. COL I, OPN, OCN, osterix and BSP), were analyzed during culture. According to the progression of bone markers during osteogenic differentiation suggested by literature, ALP activity was at its highest at early time points, and RUNX2 gene expression was upregulated from early time points, while OCN, osterix and BSP genes were highly expressed at later time points when cultured in MM. [62, 66, 67] COL I and OPN expression was maintained close to basal levels in BM, and a bit downregulated, with respect to day 7, in MM. Interestingly, no significant effect of nHA was observed in MM, suggesting that the soluble osteogenic factors might play a more pronounced role than HA in the stimulation of osteogenesis, and the absence of a synergistic effect. In the case of culture in BM, all gene expression levels were much lower compared to MM cultures, maintaining gene expression at basal levels, with a slightly upregulation at day 35. Similar to MM cultures, no major differences in the gene expression levels were observed among scaffold types when cultured in BM. Yet, immunofluorescence revealed OCN and OPN protein expression at day 35 in BM only on 45-nHA scaffolds, while the ELISA analysis showed that OCN was also secreted in BM at early timepoints only in 45-nHA scaffolds. Despite these results suggesting the potential induction of osteogenesis by 45-nHA scaffolds at late time points, this could not be concluded with a high degree of confidence, since the qualitative immunofluorescence observations were not supported by the aforementioned quantitative gene expression profiles. Moreover, OCN secretion in 45-nHA scaffolds measured by ELISA did not increase at later timepoints and levelled to other scaffold types. Previous studies have already suggested the lack of osteogenic response of cells to HA in composite scaffolds produced by ME-AM. This has been shown for polymer-HA composite scaffolds with different HA amounts. For example, PCL/PLGA-HA scaffolds with 10 wt% ceramic content in MM, as well as PEOT/PBT-HA scaffolds with 15wt% HA in BM or MM, did not enhance osteogenic differentiation of rabbit MSCs and hMSCs, respectively, when compared to pure polymer scaffolds. [39, 60] Similarly, PLA-HA scaffolds with 50 wt% HA seeded with hMSCs did not show any differences in gene expression compared to bare polymeric scaffolds when cultured in BM. [68] Consistently, our results also demonstrated that increasing the nHA-to-copolymer ratio did not enhance the osteogenic potential of the scaffolds. We hypothesize that this is due to the suboptimal ion exchange dynamics between the scaffold and the medium. On one hand, we observed that upon incubation in SPS, scaffolds were able to release Ca and P from the nHA scaffolds, at the rate of 10 to 200 μM every 3-4 days. On the other hand, ICP-MS measurements revealed overall Ca and P depletion from media, as previously reported for HA or HA-polymer composite scaffolds incubated in cell culture medium. [68–70] Therefore, it is plausible that the slow release dynamics, compared to the fast (re)precipitation events, led to insufficient Ca concentration in the medium so to affect Ca signaling pathways involved in osteogenesis. Previous reports have shown that Ca and P salts added to culture media [71–73] or released from polymeric scaffolds, [74, 75] at concentrations in the mM range, affect Ca and P signaling pathways involved in osteogenesis, leading to gene upregulation and matrix mineralization. Accordingly, scaffolds containing more soluble CaPs (e.g. BCP or TCP) and, therefore, more optimum ion exchange dynamics, have shown to outperform HA based scaffolds, both in *in vitro* and *in vivo* scenarios. [13, 69, 76, 77]

Ca and P depletion from media and SEM images of scaffolds surface after incubation in cell culture media and in SBF, confirmed CaP (re)precipitation on the scaffolds filaments surface. This was due to the presence of nHA particles on the filaments surface, negatively charged at physiological pH, acting as nucleation sites and triggering electrostatic interactions with the Ca^2+^ cations in the media, which can accumulate on the surface of scaffold forming a positively charged Ca-rich layer and stimulate phosphate anions accumulation. [78] This sequential process ultimately leads to the formation of a crystal phase of apatite, which *in vivo* has shown to promote the co-precipitation of endogenous proteins (e.g. BMPs), ultimately triggering undifferentiated cells to commit to the osteogenic lineage. [13, 40]. This information, together with the lack of osteogenic differentiation in BM, suggests that the ARS staining in 20-nHA and 45 n-HA scaffolds after 35 days of culture in BM was not an hMSCs induced effect, but rather the consequence of a mineral phase (calcium carbonate or CaP) precipitating from media onto the scaffolds’ nHA. Notably, since both 45-nHA cell seeded and non-cell seeded scaffolds depleted BM equally of Ca, and MM of Ca and P, and yet non-cell seeded scaffolds presented lower ARS staining and thinner ions layer adsorption, it is hypothesized that the ECM produced by cells on the nHA scaffolds helped to further stabilize the mineral layers, respectively. While 0-nHA scaffolds demonstrated depletion of Ca from media over the culture period, they did not show any ARS staining in BM, or CaP layer formation in SBF. This is likely due to the lack of nucleation sites for CaP precipitation, and the weaker interaction of Ca with PEOT/PBT in 0-nHA scaffolds, leading to an unstable adsorption, in contrast to the strong Ca-nHA interactions on 45-nHA scaffolds. This hypothesis is supported by previous research on PEOT/PBT copolymers calcification, showing that efficient Ca-PEO complexations only occurred in copolymers with higher PEO molecular weight and higher PEOT-PBT ratios, compared to the one used in this study. [33, 79] Consequently, it is plausible that CaP precipitated in the cell culture media.

To better elucidate the osteogenic property of the nHA containing scaffolds, longer *in vitro* culture times could be considered. Alternatively, enhancing the exposure of HA to the surface of the scaffold could potentially improve the ion exchange dynamics between scaffolds and medium and, therefore, improve the osteogenic potential of the nHA scaffolds. To do this, ME-AM scaffolds surface erosion using NaOH, [77, 80, 81] as well as bare polymeric scaffolds coating with HA by ultrasonication, [82] or pre-calcification by immersion in an SBF solution, [83] have been proposed. While these studies commonly suggest the correlation of osteogenic genes upregulation or increased matrix mineralization to the enhanced HA exposure, we believe that careful investigations are still required, as using such surface optimization methods have also shown to dramatically change the surface roughness of the scaffolds, making it hard to decouple the effect of HA bioactivity and surface roughness, as the latter is known to affect osteogenic differentiation significantly. [84]

## 5. CONCLUSION

Due to a combination of interconnected macroporosity, tunable biodegradability, optimal mechanical properties and bioactivity, 3D polymer-HA composite scaffolds prepared by ME-AM are considered important candidates, when intending to regenerate critical sized bone defects. The aim of this study was the fabrication of highly loaded PEOT/PBT-nHA composite scaffolds with up to 45 wt% nHA using ME-AM and the assessment of their mechanical and in depth biological performance as a function of the nHA content. 45-nHA scaffolds presented significantly enhanced compressive modulus, compared to bare PEOT/PBT copolymer scaffolds, which lied in the range of cancellous bone mechanical properties. In terms of cell behavior. hMSCs were able to differentiate into the osteogenic lineage in all scaffolds regardless of the HA content in MM. Although no differences were observed in osteogenic differentiation at the gene and protein level, increased matrix mineralization was observed on 45-nHA scaffolds compared to 0-nHA and 20-nHA scaffolds. While no osteogenic differentiation of cells was observed in BM, the observed ARS signal in 45-nHA scaffolds in BM suggested the precipitation of a CaP layer from Ca and P ions present in the cell culture media. Since such a CaP layer was also formed upon immersion in SBF, 45-nHA scaffolds are thought to potentially serve as osteoconductive substrates in an *in vivo* situation, favoring osteoblasts adhesion and proliferation. Overall, our results suggest the enhanced mechanical properties of the presented highly loaded composite scaffolds (45-nHA) and their ability to reprecipitate a CaP layer, thus supporting their in vivo applicability. Yet, future research and further optimization of polymer-HA composite scaffolds prepared by ME-AM for stimulating bone regeneration, as well as on the validation of their performance through in vivo studies are needed.

## Acknowledgements

We are grateful to H2020-NMP-PILOTS-2015 (GA n. 685825) for financial support. PH gratefully acknowledges the Gravitation Program ‘Materials-Driven Regeneration’, funded by the Netherlands Organization for Scientific Research (NWO) (024.003.013). Some of the materials used in this work were provided by the Texas A&M Health Science Center College of Medicine Institute for Regenerative Medicine at Scott & White through a grant from NCRR of the NIH (Grant #P40RR017447). We would also like to thank Eva Gubbins from MERLN Institute for performing the ICP-MS measurements.

## Data availability

The raw/processed data required to reproduce these findings cannot be shared at this time as the data also forms part of an ongoing study.

## Supplementary data

Supplementary data related to this article can be found at

## Notes

### Competing Interest Statement

The authors have declared no competing interest.

## References

[1] J.F. Keating, A.H. Simpson, C.M. Robinson, The management of fractures with bone loss, The Journal of bone and joint surgery. British volume 87(2) (2005) 142–50.

[2] C. Karger, T. Kishi, L. Schneider, F. Fitoussi, A.C. Masquelet, Treatment of posttraumatic bone defects by the induced membrane technique, Orthopaedics & Traumatology: Surgery & Research 98(1) (2012) 97–102.

[3] M.A. Catagni, F. Guerreschi, L. Lovisetti, Distraction osteogenesis for bone repair in the 21st century: Lessons learned, Injury 42(6) (2011) 580–586.

[4] G. Schmidmaier, R. Capanna, B. Wildemann, T. Beque, D. Lowenberg, Bone morphogenetic proteins in critical-size bone defects: what are the options?, Injury 40 (2009) S39–S43.

[5] E. Roddy, M.R. DeBaun, A. Daoud-Gray, Y.P. Yang, M.J. Gardner, Treatment of critical-sized bone defects: clinical and tissue engineering perspectives, European Journal of Orthopaedic Surgery & Traumatology 28(3) (2018) 351–362.

[6] J.R. Porter, T.T. Ruckh, K.C. Popat, Bone tissue engineering: a review in bone biomimetics and drug delivery strategies, Biotechnology progress 25(6) (2009) 1539–1560.

[7] J. Henkel, M.A. Woodruff, D.R. Epari, R. Steck, V. Glatt, I.C. Dickinson, P.F. Choong, M.A. Schuetz, D.W. Hutmacher, Bone regeneration based on tissue engineering conceptions—a 21st century perspective, Bone research 1(3) (2013) 216–248.

[8] J.-Y. Rho, L. Kuhn-Spearing, P. Zioupos, Mechanical properties and the hierarchical structure of bone, Medical Engineering & Physics 20(2) (1998) 92–102.

[9] W. Habraken, P. Habibovic, M. Epple, M. Bohner, Calcium phosphates in biomedical applications: materials for the future?, Materials Today 19(2) (2016) 69–87.

[10] S. Samavedi, A.R. Whittington, A.S. Goldstein, Calcium phosphate ceramics in bone tissue engineering: a review of properties and their influence on cell behavior, Acta biomaterialia 9(9) (2013) 8037–8045.

[11] V.P. Galván-Chacón, P. Habibovic, Deconvoluting the Bioactivity of Calcium Phosphate-Based Bone Graft Substitutes: Strategies to Understand the Role of Individual Material Properties, Advanced Healthcare Materials 6(13) (2017) 1601478.

[12] A. Barradas, H. Yuan, C.A. van Blitterswijk, P. Habibovic, Osteoinductive biomaterials: current knowledge of properties, experimental models and biological mechanisms, Eur Cell Mater 21(407) (2011) 29.

[13] P. Habibovic, H. Yuan, C.M. van der Valk, G. Meijer, C.A. van Blitterswijk, K. de Groot, 3D microenvironment as essential element for osteoinduction by biomaterials, Biomaterials 26(17) (2005) 3565–3575.

[14] D.W. Hutmacher, Scaffolds in tissue engineering bone and cartilage, Biomaterials 21(24) (2000) 2529–2543.

[15] R. Trombetta, J.A. Inzana, E.M. Schwarz, S.L. Kates, H.A. Awad, 3D Printing of Calcium Phosphate Ceramics for Bone Tissue Engineering and Drug Delivery, Annals of Biomedical Engineering 45(1) (2017) 23–44.

[16] A.P. Moreno Madrid, S.M. Vrech, M.A. Sanchez, A.P. Rodriguez, Advances in additive manufacturing for bone tissue engineering scaffolds, Materials Science and Engineering: C 100 (2019) 631–644.

[17] A. Barba, A. Diez-Escudero, Y. Maazouz, K. Rappe, M. Espanol, E.B. Montufar, M. Bonany, J.M. Sadowska, J. Guillem-Marti, C. Öhman-Mägi, C. Persson, M.C. Manzanares, J. Franch, M.P. Ginebra, Osteoinduction by Foamed and 3D-Printed Calcium Phosphate Scaffolds: Effect of Nanostructure and Pore Architecture, ACS Appl Mater Interfaces 9(48) (2017) 41722–41736.

[18] S. Raymond, Y. Maazouz, E.B. Montufar, R.A. Perez, B. González, J. Konka, J. Kaiser, M.-P. Ginebra, Accelerated hardening of nanotextured 3D-plotted self-setting calcium phosphate inks, Acta Biomaterialia 75 (2018) 451–462.

[19] A.E. Jakus, A.L. Rutz, S.W. Jordan, A. Kannan, S.M. Mitchell, C. Yun, K.D. Koube, S.C. Yoo, H.E. Whiteley, C.-P. Richter, R.D. Galiano, W.K. Hsu, S.R. Stock, E.L. Hsu, R.N. Shah, Hyperelastic “bone”: A highly versatile, growth factor–free, osteoregenerative, scalable, and surgically friendly biomaterial, Science Translational Medicine 8(358) (2016) 358ra127.

[20] M.J. Zafar, D. Zhu, Z. Zhang, 3D Printing of Bioceramics for Bone Tissue Engineering, Materials (Basel, Switzerland) 12(20) (2019).

[21] A. Kumar, S. Kargozar, F. Baino, S.S. Han, Additive Manufacturing Methods for Producing Hydroxyapatite and Hydroxyapatite-Based Composite Scaffolds: A Review, Frontiers in Materials 6(313) (2019).

[22] R.D. Bloebaum, J.G. Skedros, E.G. Vajda, K.N. Bachus, B.R. Constantz, Determining mineral content variations in bone using backscattered electron imaging, Bone 20(5) (1997) 485–90.

[23] P. Roschger, P. Fratzl, J. Eschberger, K. Klaushofer, Validation of quantitative backscattered electron imaging for the measurement of mineral density distribution in human bone biopsies, Bone 23(4) (1998) 319–26.

[24] W. Jiang, J. Shi, W. Li, K. Sun, Morphology, wettability, and mechanical properties of polycaprolactone/hydroxyapatite composite scaffolds with interconnected pore structures fabricated by a mini-deposition system, Polymer Engineering & Science 52(11) (2012) 2396–2402.

[25] W. Jiang, J. Shi, W. Li, K. Sun, Three dimensional melt-deposition of polycaprolactone/bio-derived hydroxyapatite composite into scaffold for bone repair, Journal of Biomaterials Science, Polymer Edition 24(5) (2013) 539–550.

[26] C. Esposito Corcione, F. Gervaso, F. Scalera, S.K. Padmanabhan, M. Madaghiele, F. Montagna, A. Sannino, A. Licciulli, A. Maffezzoli, Highly loaded hydroxyapatite microsphere/ PLA porous scaffolds obtained by fused deposition modelling, Ceramics International 45(2, Part B) (2019) 2803–2810.

[27] J. Yu, Y. Xu, S. Li, G.V. Seifert, M.L. Becker, Three-Dimensional Printing of Nano Hydroxyapatite/Poly(ester urea) Composite Scaffolds with Enhanced Bioactivity, Biomacromolecules 18(12) (2017) 4171–4183.

[28] J.E. Trachtenberg, J.K. Placone, B.T. Smith, J.P. Fisher, A.G. Mikos, Extrusion-based 3D printing of poly(propylene fumarate) scaffolds with hydroxyapatite gradients, J Biomater Sci Polym Ed 28(6) (2017) 532–554.

[29] T. Petrovskaya, N. Toropkov, E. Mironov, F. Azarmi, 3D printed biocompatible polylactide-hydroxyapatite based material for bone implants, Materials and Manufacturing Processes 33(16) (2018) 1899–1904.

[30] S. Chen, Y. Shi, X. Zhang, J. Ma, 3D printed hydroxyapatite composite scaffolds with enhanced mechanical properties, Ceramics International 45(8) (2019) 10991–10996.

[31] M.H. Kim, C. Yun, E.P. Chalisserry, Y.W. Lee, H.W. Kang, S.-H. Park, W.-K. Jung, J. Oh, S.Y. Nam, Quantitative analysis of the role of nanohydroxyapatite (nHA) on 3D-printed PCL/nHA composite scaffolds, Materials Letters 220 (2018) 112–115.

[32] T. Serra, J.A. Planell, M. Navarro, High-resolution PLA-based composite scaffolds via 3-D printing technology, Acta biomaterialia 9(3) (2013) 5521–5530.

[33] C. Van Blitterswijk, H. Leenders, D. Baaker, The effect of PEO ratio on degradation, calcification and bone bonding of PEO/PBT copolymer (PolyActive), Cells and Materials 3(1) (1993) 2.

[34] R. Kuijer, S.M. Bouwmeester, M.E. Drees, D.M. Surtel, E.W. Terwindt-Rouwenhorst, A. Van Der Linden, C. Van Blitterswijk, S. Bulstra, The polymer PolyactiveTM as a bone-filling substance: an experimental study in rabbits, Journal of Materials Science: Materials in Medicine 9(8) (1998) 449–455.

[35] A.A. Deschamps, M.B. Claase, W.J. Sleijster, J.D. de Bruijn, D.W. Grijpma, J. Feijen, Design of segmented poly(ether ester) materials and structures for the tissue engineering of bone, Journal of Controlled Release 78(1) (2002) 175–186.

[36] E.J.P. Jansen, J. Pieper, M.J.J. Gijbels, N.A. Guldemond, J. Riesle, L.W. Van Rhijn, S.K. Bulstra, R. Kuijer, PEOT/PBT based scaffolds with low mechanical properties improve cartilage repair tissue formation in osteochondral defects, Journal of Biomedical Materials Research Part A 89A(2) (2009) 444–452.

[37] A. Di Luca, B. Ostrowska, I. Lorenzo-Moldero, A. Lepedda, W. Swieszkowski, C. Van Blitterswijk, L. Moroni, Gradients in pore size enhance the osteogenic differentiation of human mesenchymal stromal cells in three-dimensional scaffolds, Scientific Reports 6(1) (2016) 22898.

[38] Q. Liu, J.R. De Wijn, D. Bakker, C.A. Van Blitterswijk, Surface modification of hydroxyapatite to introduce interfacial bonding with polyactiveTM 70/30 in a biodegradable composite, Journal of Materials Science: Materials in Medicine 7(9) (1996) 551–557.

[39] A. Nandakumar, C. Cruz, A. Mentink, Z.T. Birgani, L. Moroni, C. van Blitterswijk, P. Habibovic, Monolithic and assembled polymer–ceramic composites for bone regeneration, Acta biomaterialia 9(3) (2013) 5708–5717.

[40] L. Lin, K.L. Chow, Y. Leng, Study of hydroxyapatite osteoinductivity with an osteogenic differentiation of mesenchymal stem cells, Journal of Biomedical Materials Research Part A 89A(2) (2009) 326–335.

[41] R. Sinha, M. Cámara-Torres, P. Scopece, E.V. Falzacappa, A. Patelli, L. Moroni, C. Mota, A hybrid additive manufacturing platform to create bulk and surface composition gradients on scaffolds for tissue regeneration, bioRxiv (2020) 2020.06.23.165605.

[42] M. Cámara-Torres, R. Sinha, C. Mota, L. Moroni, Improving cell distribution on 3D additive manufactured scaffolds through engineered seeding media density and viscosity, Acta Biomaterialia 101 (2020) 183–195.

[43] T. Kokubo, H. Takadama, How useful is SBF in predicting in vivo bone bioactivity?, Biomaterials 27(15) (2006) 2907–2915.

[44] A.E. Jakus, A.L. Rutz, S.W. Jordan, A. Kannan, S.M. Mitchell, C. Yun, K.D. Koube, S.C. Yoo, H.E. Whiteley, C.-P. Richter, Hyperelastic “bone”: A highly versatile, growth factor–free, osteoregenerative, scalable, and surgically friendly biomaterial, Science translational medicine 8(358) (2016) 358ra127–358ra127.

[45] Y.-H. Huang, A.E. Jakus, S.W. Jordan, Z. Dumanian, K. Parker, L. Zhao, P.K. Patel, R.N. Shah, Three-dimensionally printed hyperelastic bone scaffolds accelerate bone regeneration in critical-size calvarial bone defects, Plastic and reconstructive surgery 143(5) (2019) 1397–1407.

[46] J.-T. Schantz, A. Brandwood, D.W. Hutmacher, H.L. Khor, K. Bittner, Osteogenic differentiation of mesenchymal progenitor cells in computer designed fibrin-polymer-ceramic scaffolds manufactured by fused deposition modeling, Journal of Materials Science: Materials in Medicine 16(9) (2005) 807–819.

[47] Y.F. Zhou, V. Sae-Lim, A.M. Chou, D.W. Hutmacher, T.M. Lim, Does seeding density affect in vitro mineral nodules formation in novel composite scaffolds?, Journal of Biomedical Materials Research Part A 78A(1) (2006) 183–193.

[48] L. Shor, S. Güçeri, X. Wen, M. Gandhi, W. Sun, Fabrication of three-dimensional polycaprolactone/hydroxyapatite tissue scaffolds and osteoblast-scaffold interactions in vitro, Biomaterials 28(35) (2007) 5291–5297.

[49] B. Rai, J.L. Lin, Z.X.H. Lim, R.E. Guldberg, D.W. Hutmacher, S.M. Cool, Differences between in vitro viability and differentiation and in vivo bone-forming efficacy of human mesenchymal stem cells cultured on PCL–TCP scaffolds, Biomaterials 31(31) (2010) 7960–7970.

[50] V. Karageorgiou, D. Kaplan, Porosity of 3D biomaterial scaffolds and osteogenesis, Biomaterials 26(27) (2005) 5474–5491.

[51] E.H. Backes, L.D.N. Pires, C.A.G. Beatrice, L.C. Costa, F.R. Passador, L.A. Pessan, Fabrication of Biocompatible Composites of Poly(lactic acid)/Hydroxyapatite Envisioning Medical Applications, Polymer Engineering & Science n/a(n/a) (2020).

[52] Y. Huang, G. Zhou, L. Zheng, H. Liu, X. Niu, Y. Fan, Micro-/nano-sized hydroxyapatite directs differentiation of rat bone marrow derived mesenchymal stem cells towards an osteoblast lineage, Nanoscale 4(7) (2012) 2484–90.

[53] X. Yang, Y. Li, X. Liu, R. Zhang, Q. Feng, In Vitro Uptake of Hydroxyapatite Nanoparticles and Their Effect on Osteogenic Differentiation of Human Mesenchymal Stem Cells, Stem cells international 2018 (2018) 2036176.

[54] T.M. Keaveny, E.F. Morgan, O.C. Yeh, Bone mechanics, Standard handbook of biomedical engineering and design (2004) 1–24.

[55] D.T. Reilly, A.H. Burstein, The elastic and ultimate properties of compact bone tissue, J Biomech 8(6) (1975) 393–405.

[56] T. Albrektsson, C. Johansson, Osteoinduction, osteoconduction and osseointegration, European spine journal 10(2) (2001) S96–S101.

[57] P. Habibovic, T. Woodfield, K. de Groot, C. van Blitterswijk, Predictive value of in vitro and in vivo assays in bone and cartilage repair--what do they really tell us about the clinical performance?, Advances in experimental medicine and biology 585 (2006) 327–60.

[58] S. Noda, N. Kawashima, M. Yamamoto, K. Hashimoto, K. Nara, I. Sekiya, T. Okiji, Effect of cell culture density on dental pulp-derived mesenchymal stem cells with reference to osteogenic differentiation, Scientific Reports 9(1) (2019) 5430.

[59] C.E. Holy, M.S. Shoichet, J.E. Davies, Engineering three‐dimensional bone tissue in vitro using biodegradable scaffolds: Investigating initial cell‐seeding density and culture period, Journal of Biomedical Materials Research: An Official Journal of The Society for Biomaterials, The Japanese Society for Biomaterials, and The Australian Society for Biomaterials and the Korean Society for Biomaterials 51(3) (2000) 376–382.

[60] K.K. Moncal, D.N. Heo, K.P. Godzik, D.M. Sosnoski, O.D. Mrowczynski, E. Rizk, V. Ozbolat, S.M. Tucker, E.M. Gerhard, M. Dey, G.S. Lewis, J. Yang, I.T. Ozbolat, 3D printing of poly(ε-caprolactone)/poly(D,L-lactide-co-glycolide)/hydroxyapatite composite constructs for bone tissue engineering, Journal of Materials Research 33(14) (2018) 1972–1986.

[61] B. Huang, G. Caetano, C. Vyas, J.J. Blaker, C. Diver, P. Bártolo, Polymer-Ceramic Composite Scaffolds: The Effect of Hydroxyapatite and β-tri-Calcium Phosphate, Materials 11(1) (2018) 129.

[62] T.A. Owen, M. Aronow, V. Shalhoub, L.M. Barone, L. Wilming, M.S. Tassinari, M.B. Kennedy, S. Pockwinse, J.B. Lian, G.S. Stein, Progressive development of the rat osteoblast phenotype in vitro: reciprocal relationships in expression of genes associated with osteoblast proliferation and differentiation during formation of the bone extracellular matrix, Journal of cellular physiology 143(3) (1990) 420–30.

[63] K.K. Ivaska, T.A. Hentunen, J. Vaaraniemi, H. Ylipahkala, K. Pettersson, H.K. Vaananen, Release of intact and fragmented osteocalcin molecules from bone matrix during bone resorption in vitro, The Journal of biological chemistry 279(18) (2004) 18361–9.

[64] T. Komori, H. Yagi, S. Nomura, A. Yamaguchi, K. Sasaki, K. Deguchi, Y. Shimizu, R.T. Bronson, Y.H. Gao, M. Inada, M. Sato, R. Okamoto, Y. Kitamura, S. Yoshiki, T. Kishimoto, Targeted disruption of Cbfa1 results in a complete lack of bone formation owing to maturational arrest of osteoblasts, Cell 89(5) (1997) 755–64.

[65] Y. Li, C. Ge, J.P. Long, D.L. Begun, J.A. Rodriguez, S.A. Goldstein, R.T. Franceschi, Biomechanical stimulation of osteoblast gene expression requires phosphorylation of the RUNX2 transcription factor, Journal of bone and mineral research: the official journal of the American Society for Bone and Mineral Research 27(6) (2012) 1263–74.

[66] H.I. Roach, WHY DOES BONE MATRIX CONTAIN NON-COLLAGENOUS PROTEINS? THE POSSIBLE ROLES OF OSTEOCALCIN, OSTEONECTIN, OSTEOPONTIN AND BONE SIALOPROTEIN IN BONE MINERALISATION AND RESORPTION, Cell Biology International 18(6) (1994) 617–628.

[67] R. Ozawa, Y. Yamada, T. Nagasaka, M. Ueda, A comparison of osteogenesis-related gene expression of mesenchymal stem cells during the osteoblastic differentiation induced by type-I collagen and/or fibronectin, International Journal of Oral-Medical Sciences 1(2) (2003) 139–146.

[68] L. Sun, C.B. Danoux, Q. Wang, D. Pereira, D. Barata, J. Zhang, V. LaPointe, R. Truckenmüller, C. Bao, X. Xu, P. Habibovic, Independent effects of the chemical and microstructural surface properties of polymer/ceramic composites on proliferation and osteogenic differentiation of human MSCs, Acta Biomaterialia 42 (2016) 364–377.

[69] A.M. Barradas, V. Monticone, M. Hulsman, C. Danoux, H. Fernandes, Z. Tahmasebi Birgani, F. Barrere-de Groot, H. Yuan, M. Reinders, P. Habibovic, C. van Blitterswijk, J. de Boer, Molecular mechanisms of biomaterial-driven osteogenic differentiation in human mesenchymal stromal cells, Integrative biology: quantitative biosciences from nano to macro 5(7) (2013) 920–31.

[70] Z. Tahmasebi Birgani, C.A. van Blitterswijk, P. Habibovic, Monolithic calcium phosphate/poly(lactic acid) composite versus calcium phosphate-coated poly(lactic acid) for support of osteogenic differentiation of human mesenchymal stromal cells, Journal of Materials Science: Materials in Medicine 27(3) (2016) 54.

[71] A.M.C. Barradas, H.A.M. Fernandes, N. Groen, Y.C. Chai, J. Schrooten, J. van de Peppel, J.P.T.M. van Leeuwen, C.A. van Blitterswijk, J. de Boer, A calcium-induced signaling cascade leading to osteogenic differentiation of human bone marrow-derived mesenchymal stromal cells, Biomaterials 33(11) (2012) 3205–3215.

[72] Y.C. Chai, S.J. Roberts, J. Schrooten, F.P. Luyten, Probing the Osteoinductive Effect of Calcium Phosphate by Using an In Vitro Biomimetic Model, Tissue Engineering Part A 17(7-8) (2010) 1083–1097.

[73] G.R. Beck, B. Zerler, E. Moran, Phosphate is a specific signal for induction of osteopontin gene expression, Proceedings of the National Academy of Sciences 97(15) (2000) 8352–8357.

[74] P. Habibovic, D.C. Bassett, C.J. Doillon, C. Gerard, M.D. McKee, J.E. Barralet, Collagen Biomineralization In Vivo by Sustained Release of Inorganic Phosphate Ions, Advanced Materials 22(16) (2010) 1858–1862.

[75] C.B.S.S. Danoux, D.C. Bassett, Z. Othman, A.I. Rodrigues, R.L. Reis, J.E. Barralet, C.A. van Blitterswijk, P. Habibovic, Elucidating the individual effects of calcium and phosphate ions on hMSCs by using composite materials, Acta Biomaterialia 17 (2015) 1–15.

[76] H. Yuan, H. Fernandes, P. Habibovic, J. de Boer, A.M.C. Barradas, A. de Ruiter, W.R. Walsh, C.A. van Blitterswijk, J.D. de Bruijn, Osteoinductive ceramics as a synthetic alternative to autologous bone grafting, Proc Natl Acad Sci U S A 107(31) (2010) 13614–13619.

[77] E. Nyberg, A. Rindone, A. Dorafshar, W.L. Grayson, Comparison of 3D-Printed Poly-☐-Caprolactone Scaffolds Functionalized with Tricalcium Phosphate, Hydroxyapatite, Bio-Oss, or Decellularized Bone Matrix, Tissue Engineering Part A 23(11-12) (2016) 503–514.

[78] H.-M. Kim, T. Himeno, M. Kawashita, T. Kokubo, T. Nakamura, The mechanism of biomineralization of bone-like apatite on synthetic hydroxyapatite: an in vitro assessment, Journal of The Royal Society Interface 1(1) (2004) 17–22.

[79] R. Thoma, T. Hung, E. Nyilas, A. Haubold, R. Phillips, Metal ion complexation of poly (ether) urethanes, Advances in biomedical polymers, Springer1987, pp. 131–145.

[80] Y.S. Cho, S. Choi, S.-H. Lee, K.K. Kim, Y.-S. Cho, Assessments of polycaprolactone/hydroxyapatite composite scaffold with enhanced biomimetic mineralization by exposure to hydroxyapatite via a 3D-printing system and alkaline erosion, European Polymer Journal 113 (2019) 340–348.

[81] Y.S. Cho, M. Quan, S.-H. Lee, M.W. Hong, Y.Y. Kim, Y.-S. Cho, Assessment of osteogenesis for 3D-printed polycaprolactone/hydroxyapatite composite scaffold with enhanced exposure of hydroxyapatite using rat calvarial defect model, Composites Science and Technology 184 (2019) 107844.

[82] J. Si, J. Lin, C. Su, S. Yu, Z. Cui, Q. Wang, W. Chen, L.-S. Turng, Ultrasonication-Induced Modification of Hydroxyapatite Nanoparticles onto a 3D Porous Poly(lactic acid) Scaffold with Improved Mechanical Properties and Biocompatibility, Macromolecular Materials and Engineering 304(7) (2019) 1900081.

[83] A. Nandakumar, A. Barradas, J. de Boer, L. Moroni, C. van Blitterswijk, P. Habibovic, Combining technologies to create bioactive hybrid scaffolds for bone tissue engineering, Biomatter 3(2) (2013) e23705.

[84] A.B. Faia-Torres, S. Guimond-Lischer, M. Rottmar, M. Charnley, T. Goren, K. Maniura-Weber, N.D. Spencer, R.L. Reis, M. Textor, N.M. Neves, Differential regulation of osteogenic differentiation of stem cells on surface roughness gradients, Biomaterials 35(33) (2014) 9023–9032.

