## Supplementary Information for "Effect of Highly Loaded Nanohydroxyapatite Composite Scaffolds Prepared via Melt Extrusion Additive Manufacturing on the Osteogenic Differentiation of Human Mesenchymal Stromal Cells"

M. Cámara-Torres, Dr. R. Sinha, Prof. P. Habibovic, Dr. C. Mota, Prof. L. Moroni

Maastricht University, MERLN Institute for Technology-Inspired Regenerative Medicine, Complex Tissue regeneration Department, 6229 ER Maastricht, The Netherlands.

Dr. A. Sánchez

TECNALIA, Basque Research and Technology Alliance (BRTA), 20009 Donostia-San Sebastian, Spain.

Prof. A. Patelli

Department of Physics and Astronomy, Padova University, 35131 Padova, Italy.

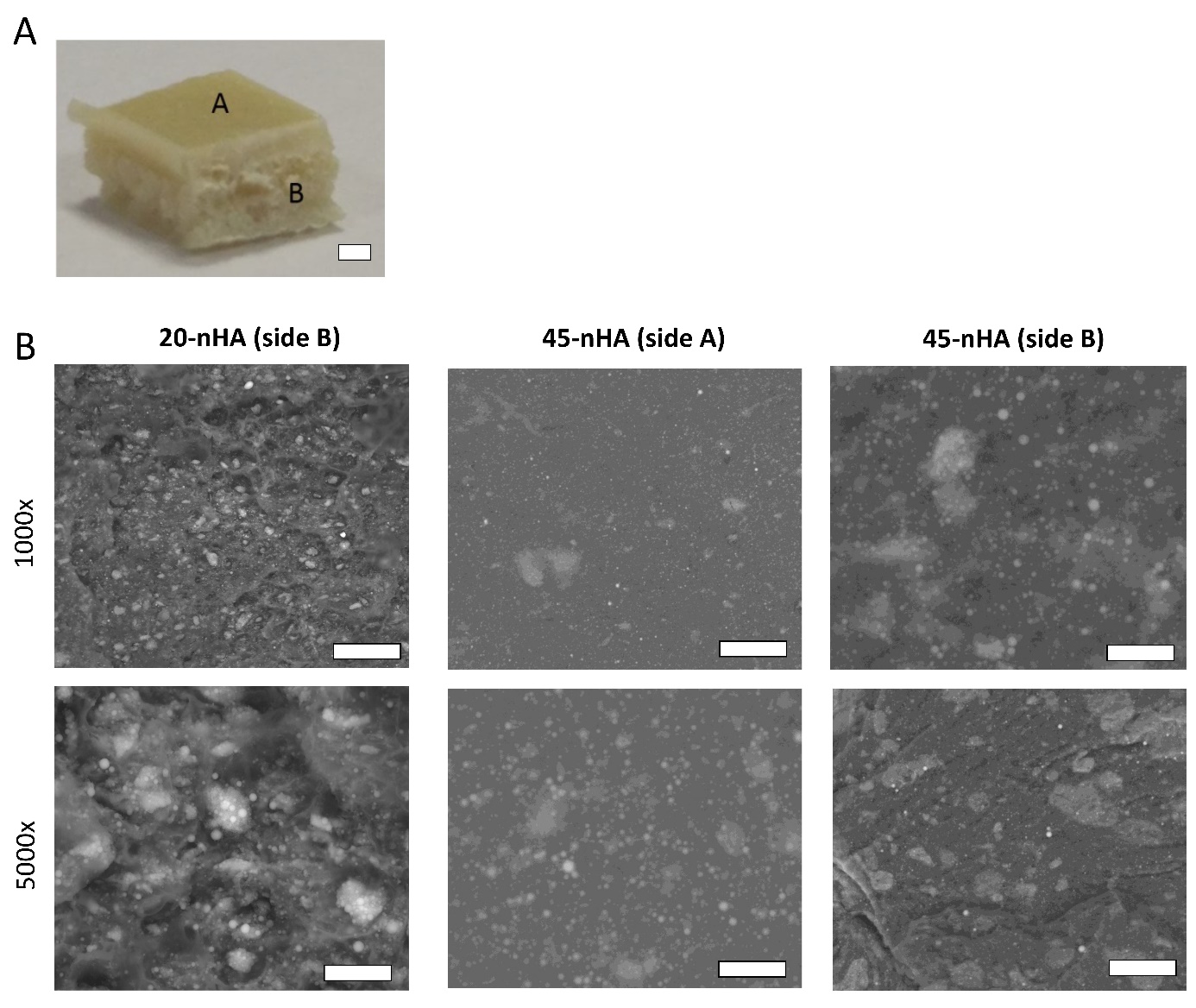

**Figure S1**. (A) Image of a 45-nHA pellet obtained after the solvent blending of nHA and PEOT/PBT process. The top side of the pellets is marked as side A and the cross section of the pellet as side B. Scale bar: 1 mm. (B) BSEM micrographs of the side B of 20-nHA pellets, and the side A and B of 45-nHA pellets at two different magnifications (1000x and 5000x). Scale bars top row 50 µm, lower row 10 µm.

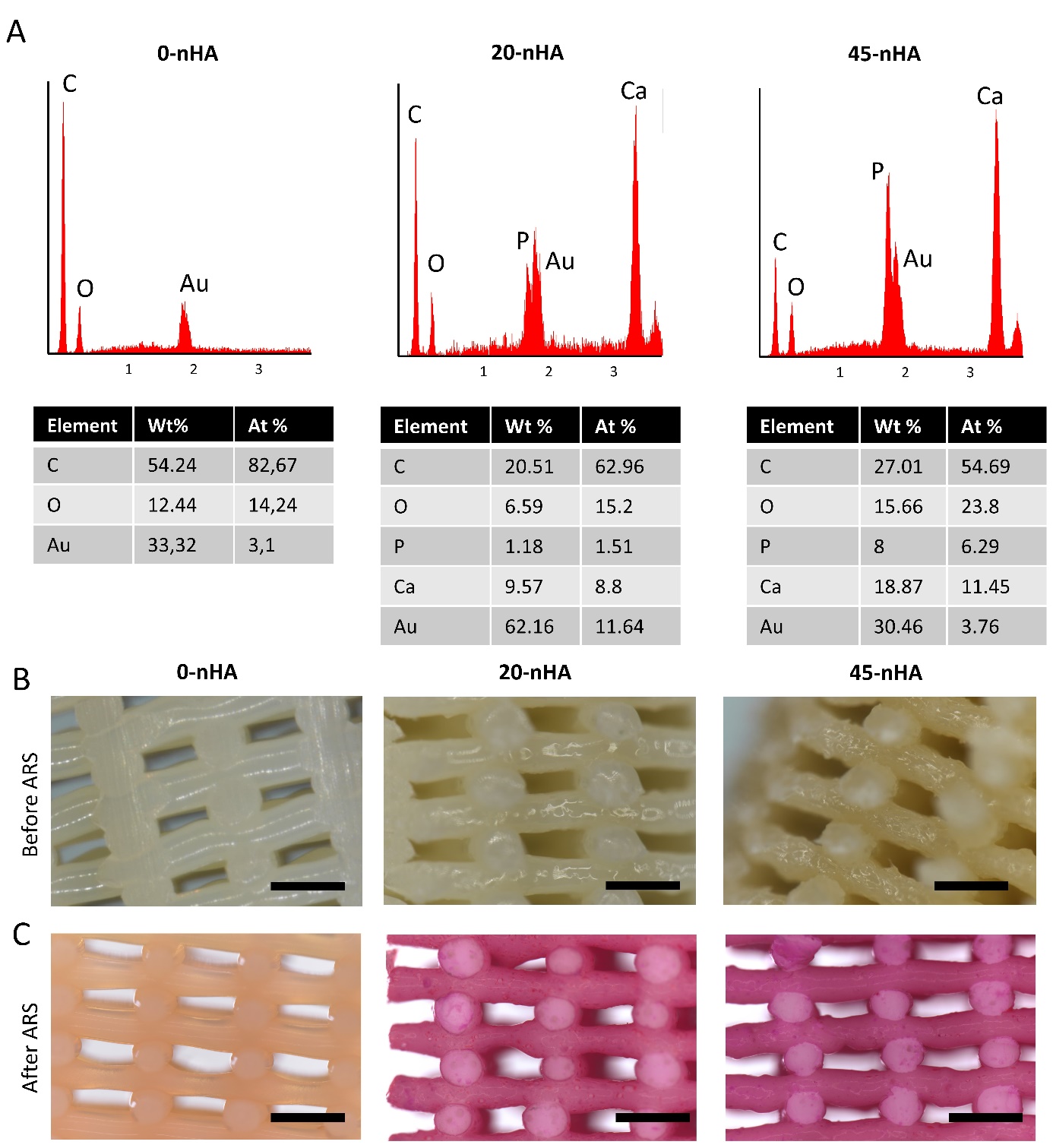

**Figure S2.** (A) Elemental analysis spectrum obtained with EDS and corresponding elements weight and atomic % of a ~80 x 80 µm^2^ area from a 0-nHA and 45-nHA surface filament and 45-nHA cross section. Stereomicroscopy images of a- prepared 0-nHA, 20-nHA and 45-nHA scaffolds (A) before alizarin red (ARS) staining and (B) after ARS staining.

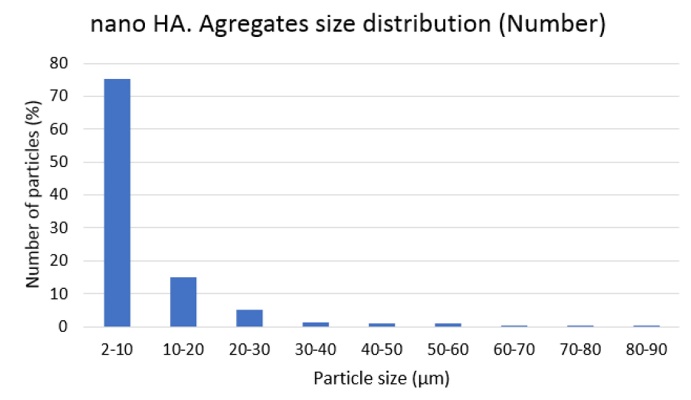

**Figure S3.** nHA aggregates size distribution on 45-nHA scaffolds.

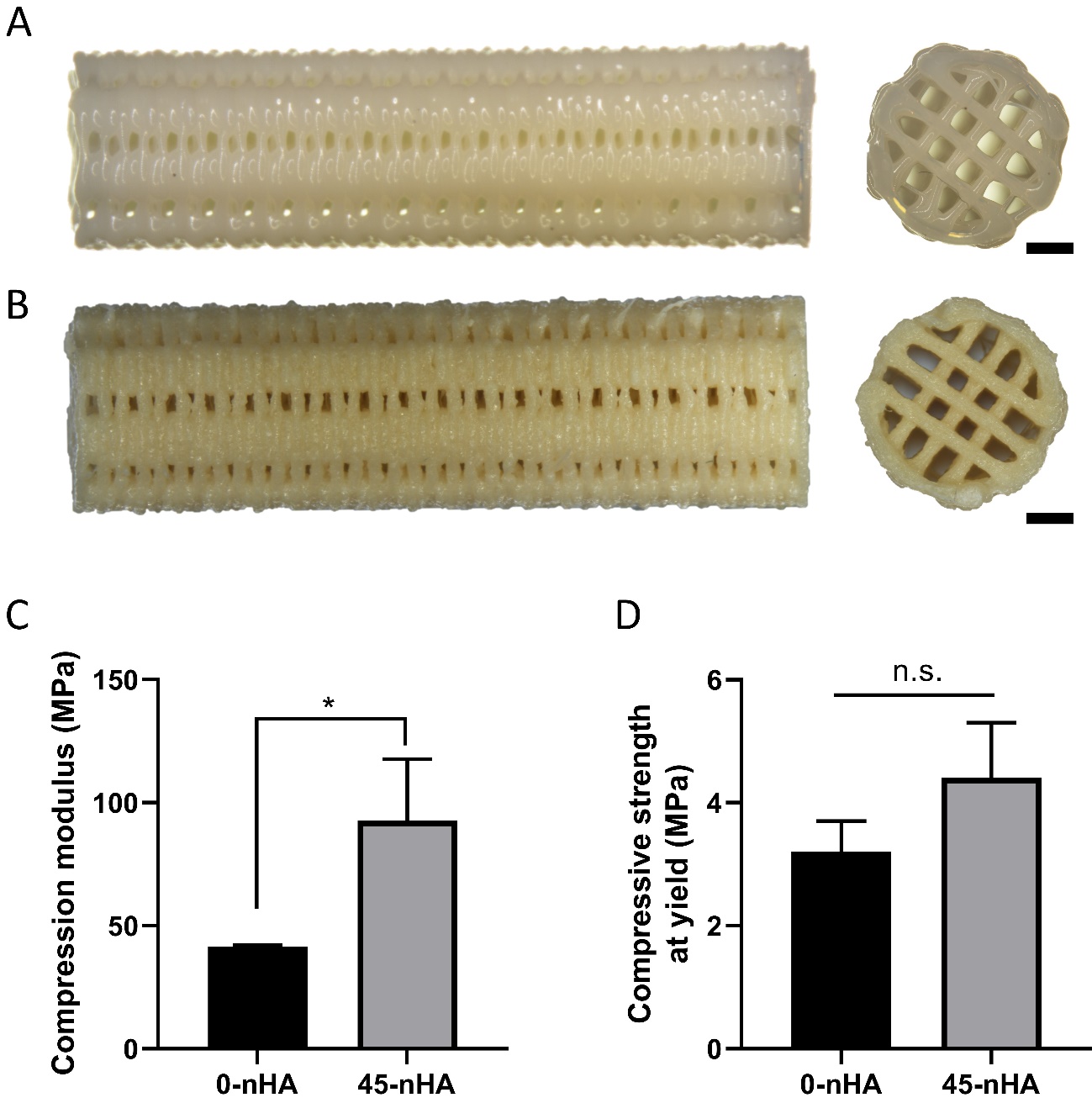

**Figure S4.** (A) Cylindrical 0-nHA and (B) 45-nHA scaffolds with 15 mm height and 4 mm diameter. Scale bars 1 mm. (C, D) Mechanical properties in compression of 0-nHA and 45-nHA scaffolds.

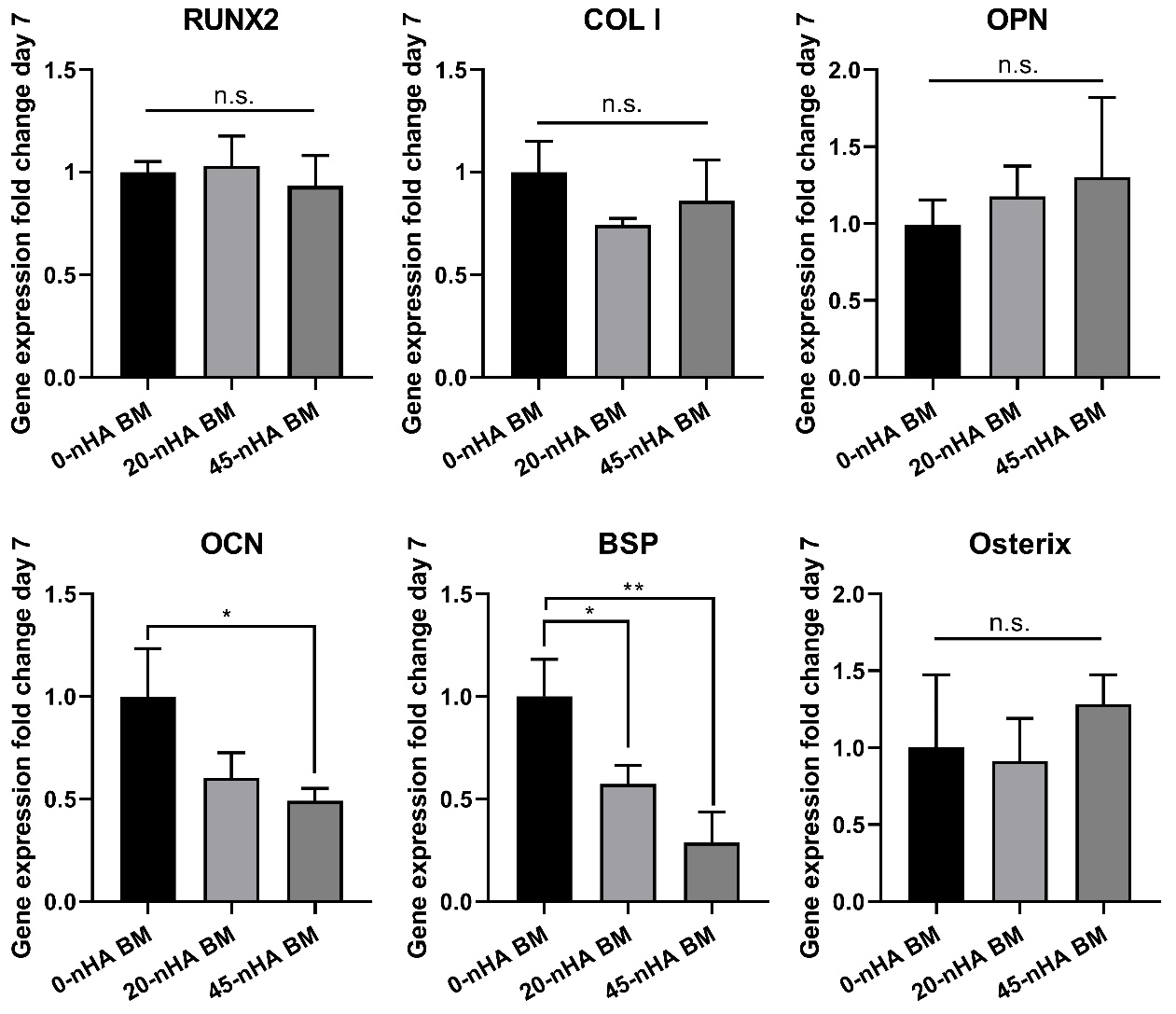

**Figure S5.** Gene expression fold change of hMSCs cultured on scaffolds with different nHA concentrations for 7 days in BM. Normalized to 0-nHA d7 BM. Data presented as average ± s.d., and statistical significance performed using one-way ANOVA with Tukey’s multiple comparison test (n.s. p>0.05; *$ p < 0.05; **$$ p < 0.01)

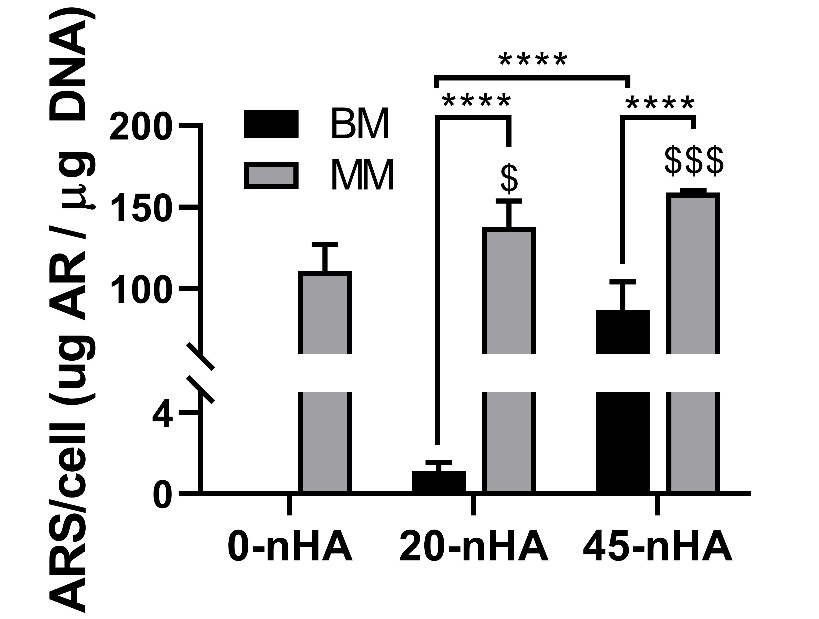

**Figure S6.** Quantification of the alizarin red S staining extracted from scaffolds after 35 days of culture in BM or MM normalized to cell number. Data presented as average ± s.d. and statistical significance performed using two-way ANOVA with Tukey’s multiple comparison test. *$ p < 0.05; ***$$$ p < 0.001; ****$$$$ p < 0.0001; * for comparisons among BM and MM each scaffold type; $ for comparisons among scaffold types each culture media type.

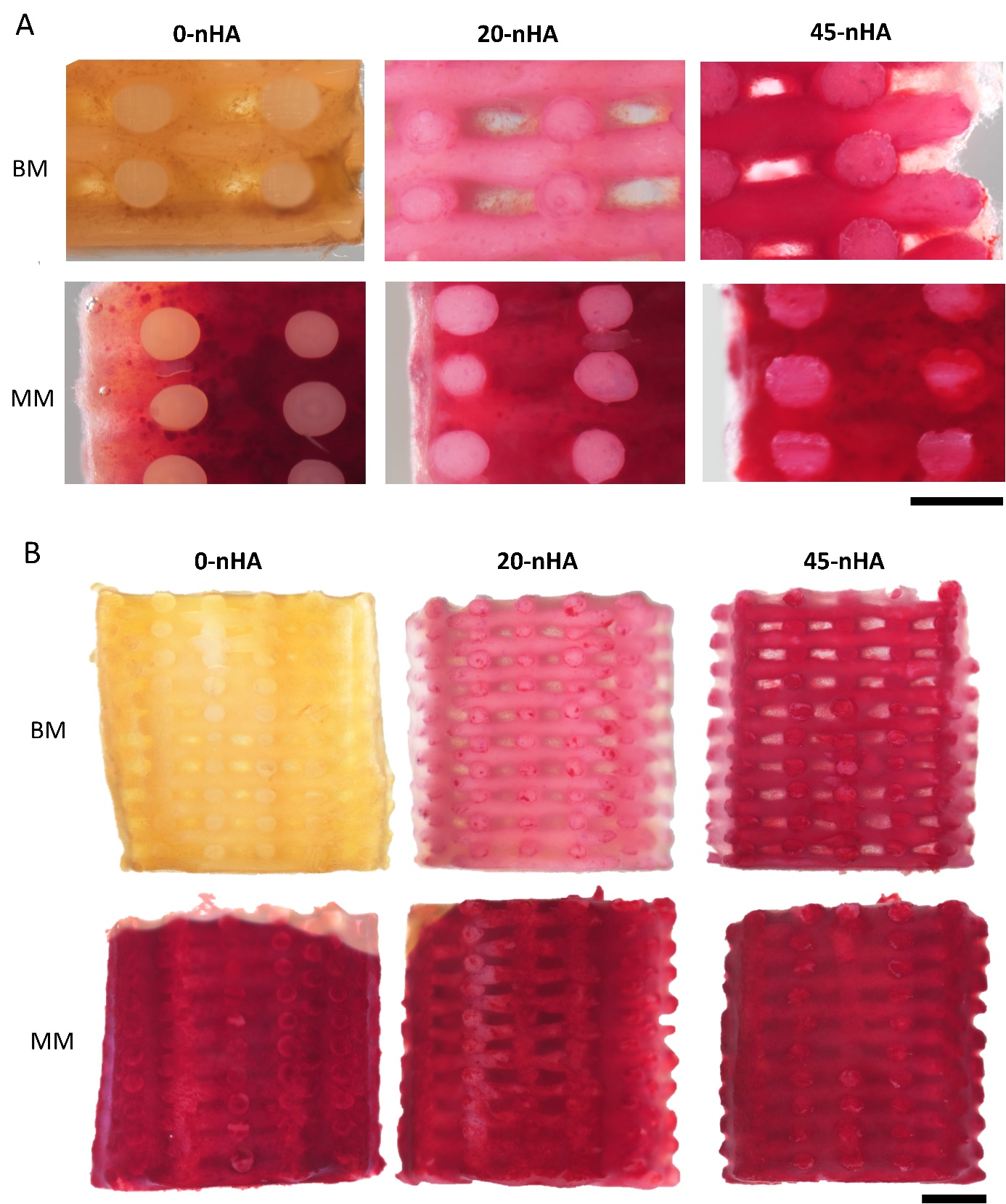

**Figure S7.** Stereomicroscopy images of scaffolds stained with ARS after 35 days of culture in BM or MM (7 days in BM followed by 28 days in MM). (A) High magnification images of scaffolds cross sections. Scale bar 500 µm. (B) Images of scaffolds outer surface of the scaffolds. Scale bar: 1 mm.

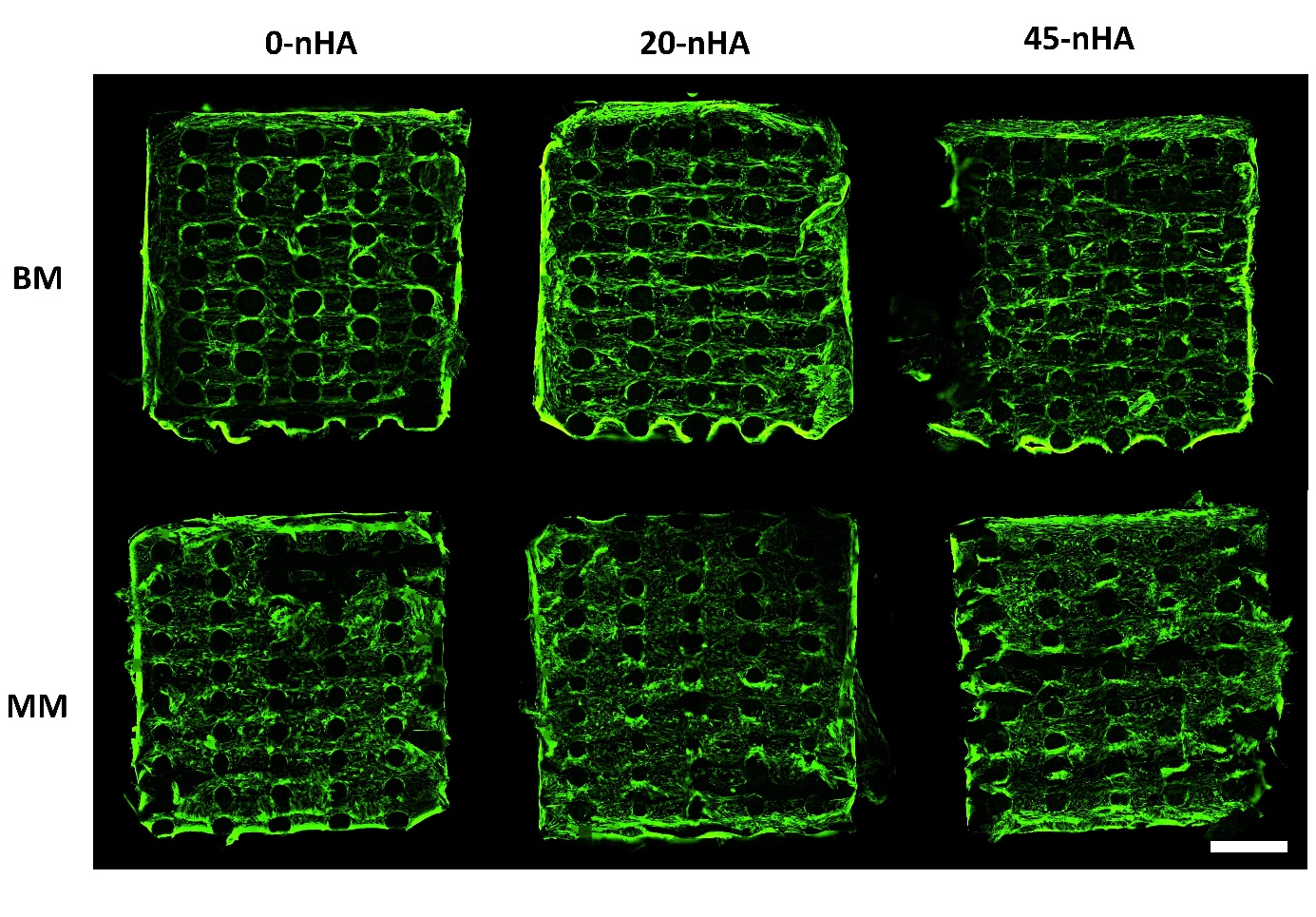

**Figure S8.** Fluorescent images of hMSCs (F-actin, green) on scaffolds cross sections after 35 days of culture in BM or MM. Scale bar: 1 mm.

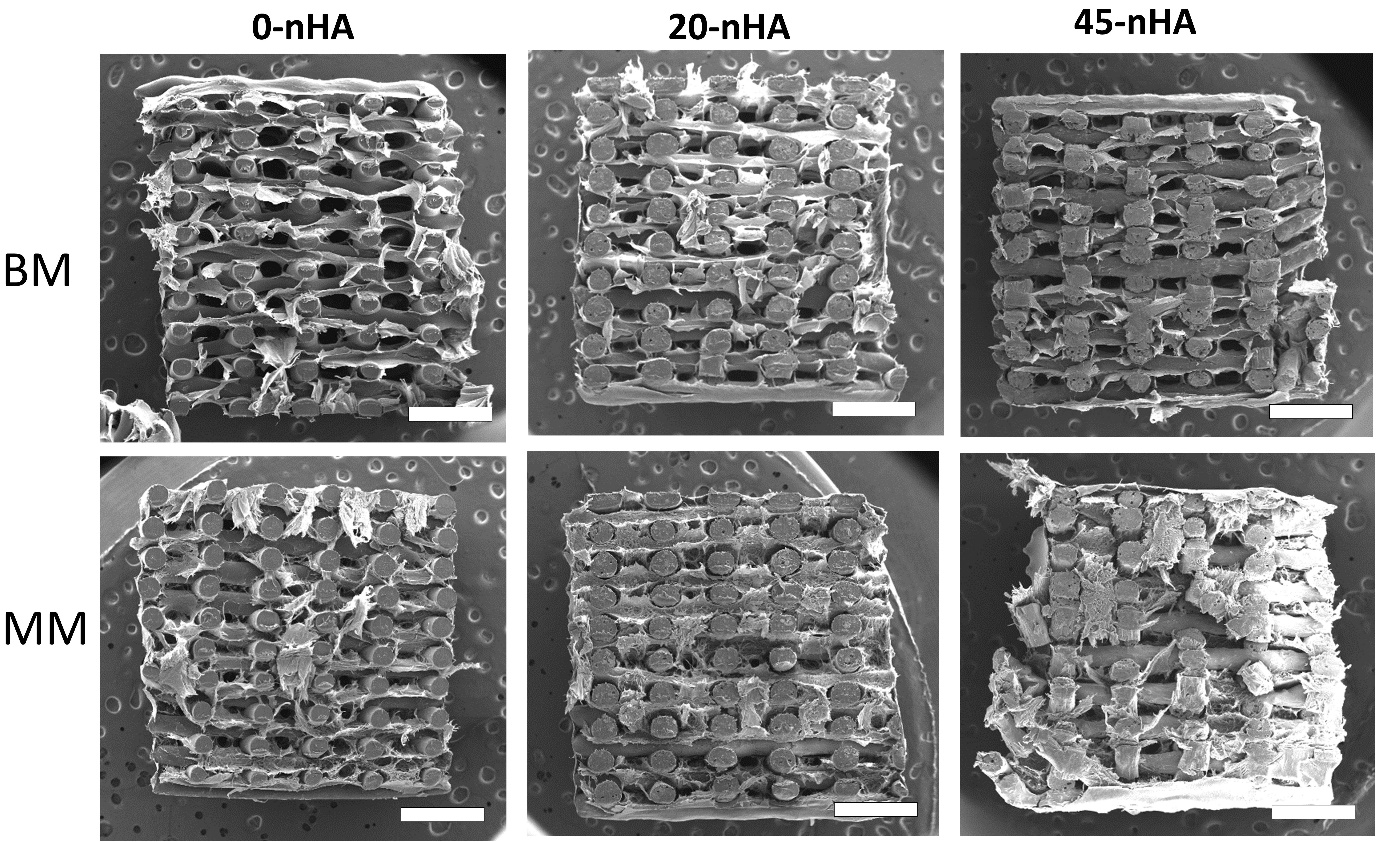

**Figure S9.** SEM micrographs of scaffolds cross sections after 35 days of culture in BM or MM. Scale bars: 1 mm.

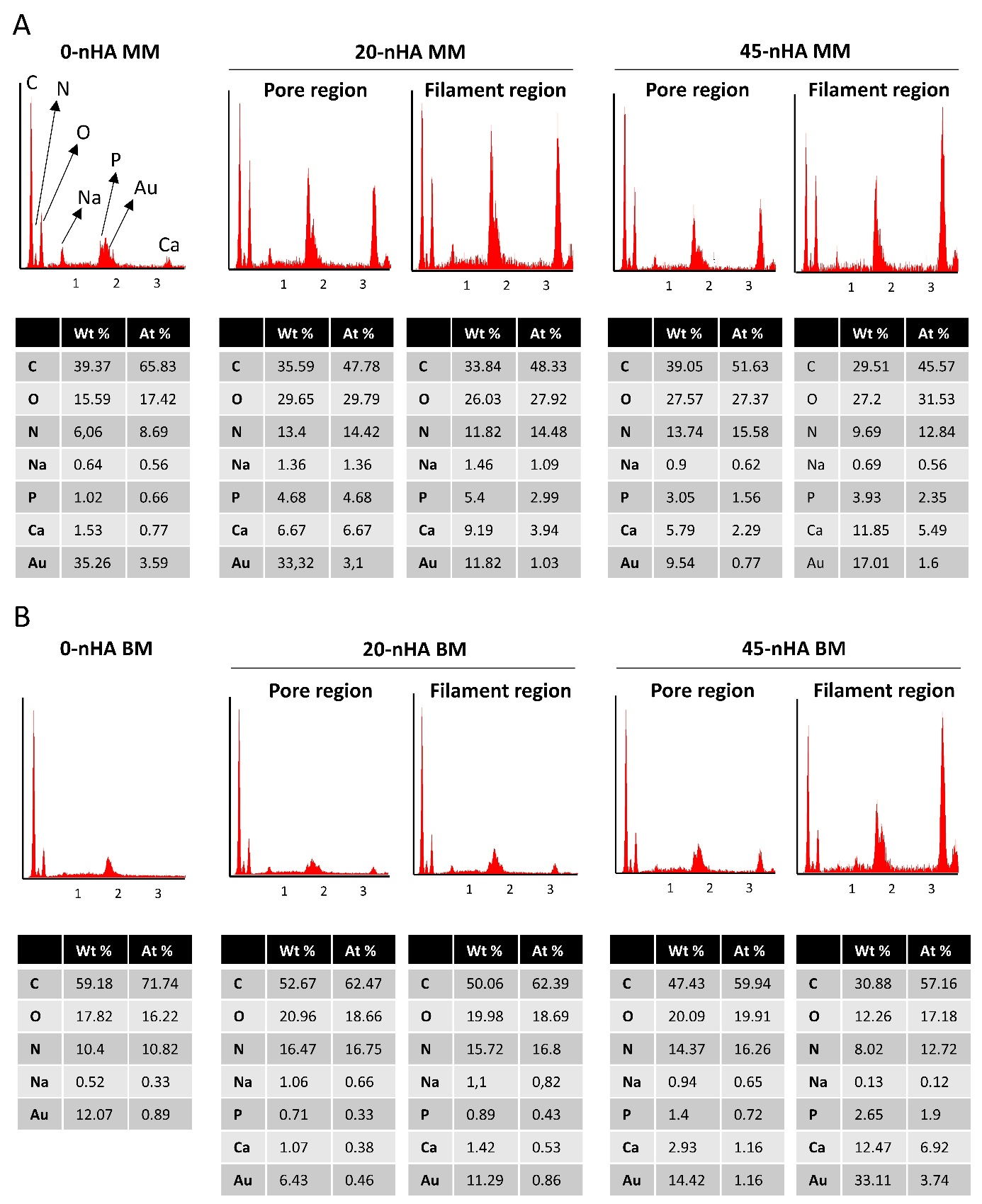

**Figure S10.** Elemental analysis spectrum obtained with EDS and corresponding elements weight and atomic % of a ~80 x 80 µm2 area of the ECM formed in 0-nHA and 45-nHA scaffolds after 35 days in culture in BM or MM

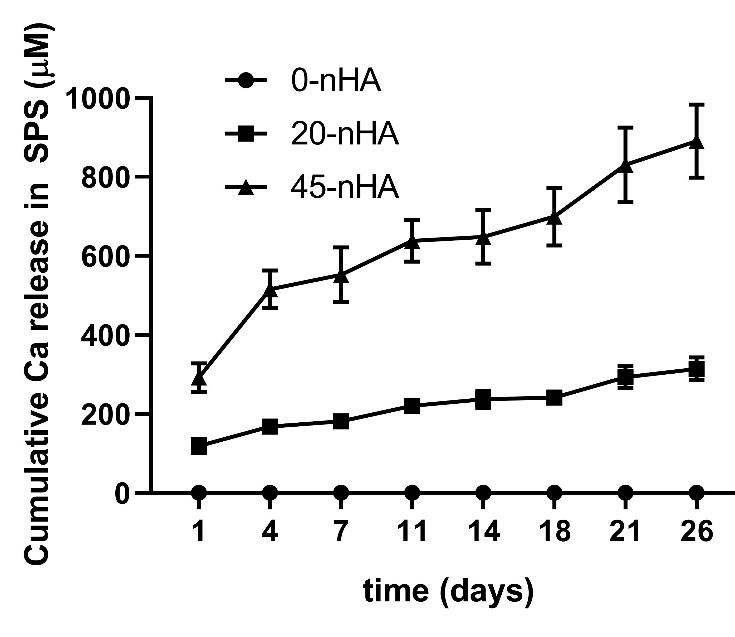

**Figure S11.** Ca ions release over time from nHA composite scaffolds upon immersion in SPS, measured by ICP-MS.

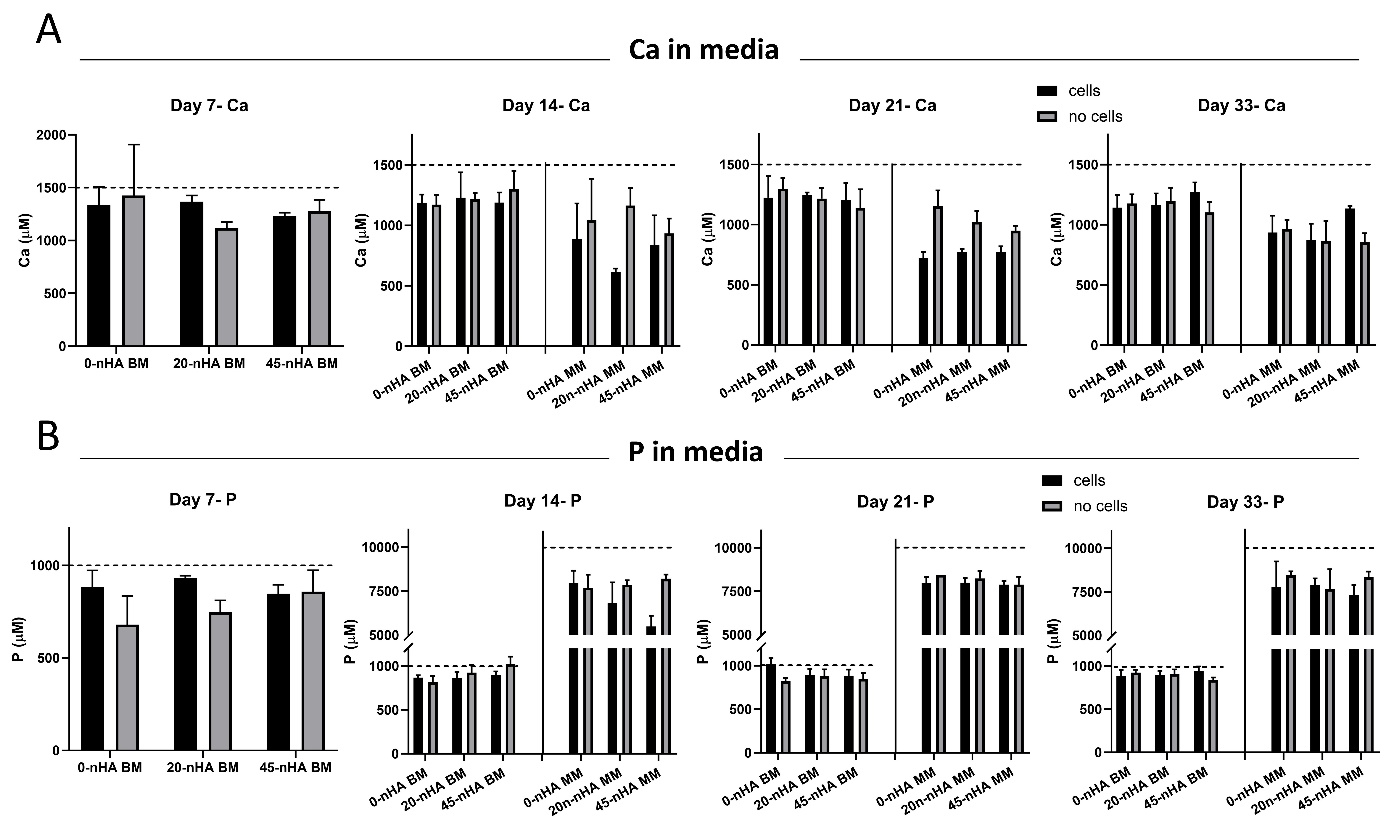

**Figure S12.** ICP-MS measurements of the calcium and phosphorous concentrations in cell culture medium after 7, 14, 21, and 35 days of culture on 0-nHA, 20-nHA and 45-nHA scaffolds with and without cells. The dash lines represent fresh medium concentrations. Data presented as average ± s.d.

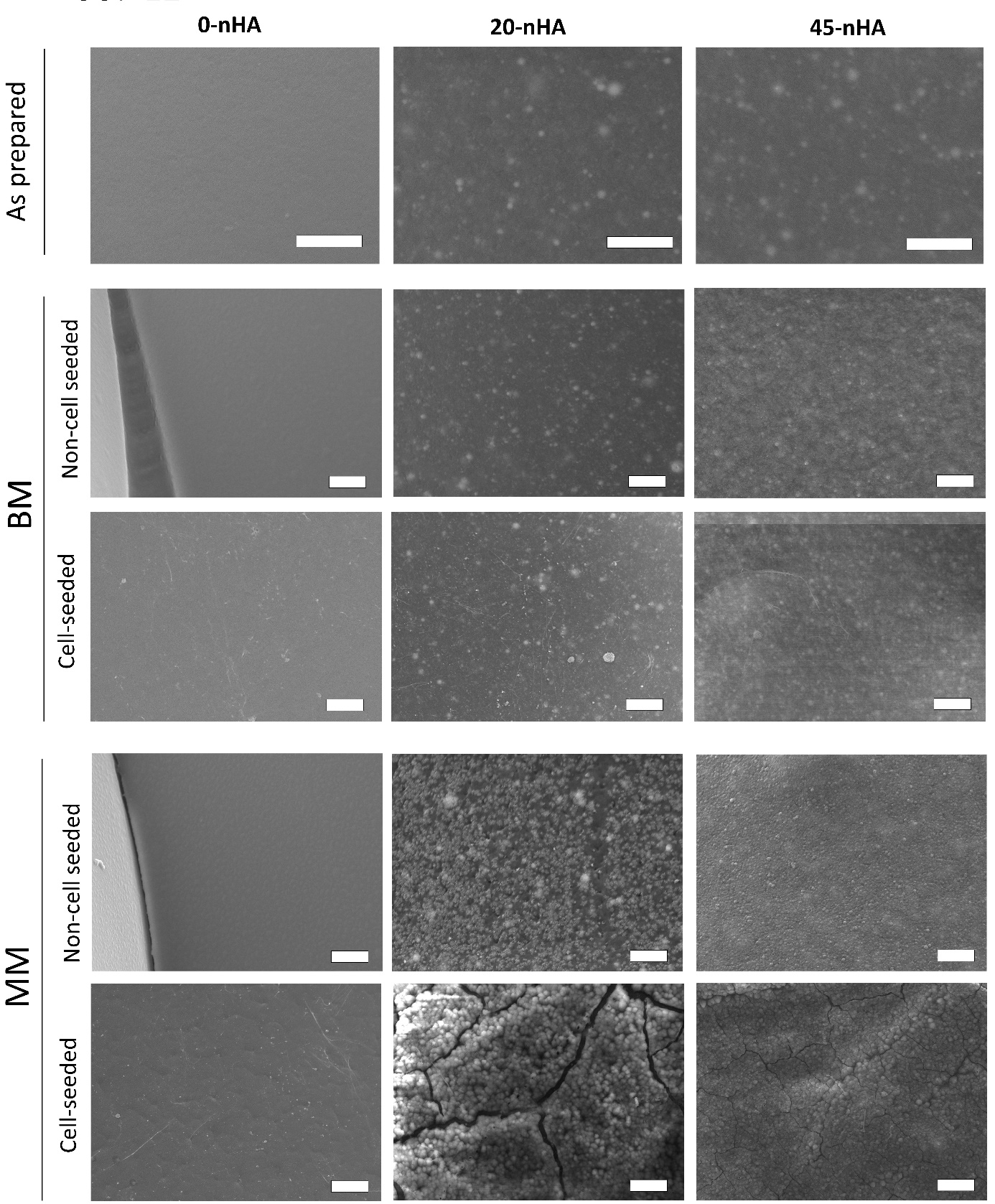

**Figure S13.** Representative SEM micrographs of 0-nHA, 20-nHA and 45-nHA non-cell seeded and cell seeded scaffolds incubated for 35 days in BM or MM. Scale bars: 20 µm.

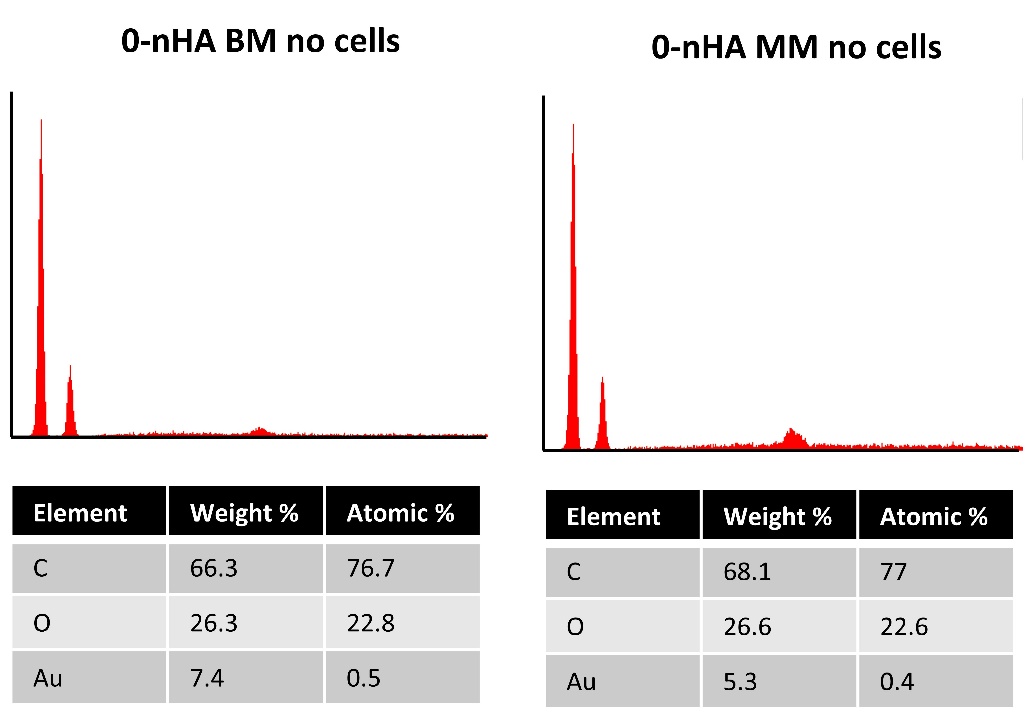

**Figure S14.** Elemental analysis spectrum obtained with EDS and corresponding elements weight and atomic % of a ~80 x 80 µm2 area of the surface of filaments in 0-nHA non-cell seeded scaffolds after 35 days in culture in BM or MM.
